# microRNA-dependent regulation of biomechanical genes establishes tissue stiffness homeostasis

**DOI:** 10.1101/359521

**Authors:** Albertomaria Moro, Tristan Discroll, William Armero, Liana C. Boraas, Dionna M. Kasper, Nicolas Baeyens, Charlene Jouy, Venkatesh Mallikarjun, Joe Swift, Sang Joon Ahn, Donghoon Lee, Jing Zhang, Mengting Gu, Mark Gerstein, Martin Schwart, Stefania Nicoli

## Abstract

The mechanical properties of tissues, which are determined primarily by their extracellular matrix (ECM), are largely stable over time despite continual turnover of ECM constituents ^1,2^. These observations imply active homeostasis, where cells sense and adjust rates of matrix synthesis, assembly and degradation to keep matrix and tissue properties within the optimal range. However, the regulatory pathways that mediate this process are essentially unknown^3^. Genome-wide analyses of endothelial cells revealed abundant microRNA-mediated regulation of cytoskeletal, adhesive and extracellular matrix (CAM) mRNAs. High-throughput assays showed co-transcriptional regulation of microRNA and CAM genes on stiff substrates, which buffers CAM expression. Disruption of global or individual microRNA-dependent suppression of CAM genes induced hyper-adhesive, hyper-contractile phenotypes in multiple systems *in vitro,* and increased tissue stiffness in the zebrafish fin-fold during homeostasis and regeneration *in vivo.* Thus, a network of microRNAs and CAM mRNAs mediate tissue mechanical homeostasis.

---

Cells sense physical forces, including the stiffness of their ECM, through mechanosensitive integrins, their associated proteins, and actomyosin. These factors transduce physical forces into biochemical signals that regulate gene expression and cell function^2,3^. Tissues maintain nearly constant physical properties in the face of growth, injury, ECM turnover, and altered external forces (e.g. from blood pressure, tissue hydration or body weight)^1,4,5^. These effects imply tissue mechanical homeostasis, in which cells sense mechanical loads, due to both external and internal forces, and adjust their rates of matrix synthesis, degradation and organization to keep tissue properties constant. Cell contractility is critical in this process, as it is a key component of both the stiffness-sensing regulatory pathways and of the matrix assembly process that governs resultant matrix properties, including stiffness^2,6^.

Mechanical homeostasis requires that integrin mechanotransduction pathways mediate negative feedback regulation of the contractile and biosynthetic pathways to maintain optimal tissue stiffness. That is, too soft/low force triggers increased matrix synthesis and contractility, while too stiff/high force triggers the opposite. However, *in vitro* studies have mainly elucidated positive feedback (or feed forward) circuits, where rigid substrates or high external forces increase actin myosin contraction, focal adhesions and ECM synthesis^7^. This type of mechanotransduction signaling often characterizes fibrotic tissues, where sustained contractility and excessive ECM compromise tissue function. Very little is known about negative feedback pathways that are therefore critical to establish proper stiffness/contractility in normal, healthy tissues.

MicroRNAs (miRNAs) regulate gene expression at the post-transcriptional level ^8-10^, and often function to reduce fluctuations in protein levels caused by changes in transcriptional inputs or extracellular factors. miRNAs therefore participate in regulatory feedback loops that contribute to homeostasis in multiple contexts ^11-13^. Studies of miRNA regulation of biological processes often focus on a single or a few miRNA-target gene interactions^14-18^. However, miRNAs appear to function within larger networks that are likely critical for cellular functions^11,19^.

miRNAs regulate target mRNAs via homologous base pairing. After transcription, miRNAs are processed via the ribonucleases DROSHA/DRG8 and DICER^20^ into mature 20-21 nucleotide (nt) hairpins that recognize abundant and conserved 7-8 nt miRNA responsive elements (MREs) within mRNAs. MREs reside mainly in the 3’ untranslated regions (3’UTR) of mRNAs and base-pair with the 5’ miRNA mature sequence (SEED region)^21^. The miRNA-MRE pairs are recognized by the Argonaute2 (AGO2) protein complex, resulting in mRNA destabilization and/or reduced protein expression^20^.

To investigate a potential role for miRNAs in mechanical homeostasis, we analyzed miRNA-mRNA interactions transcriptome-wide using an AGO2-HITS-CLIP approach^22^. AGO2-bound miRNAs/mRNAs were isolated from two unrelated human endothelial cells (EC) lines, which are known to respond to mechanical forces, including ECM loads^3,23^ We exposed cultured human umbilical artery ECs (HUAECs) and human venous umbilical ECs (HUVECs) to UV light to cross-link protein-RNA complexes. Subsequently, we immunoprecipitated AGO2-RNA complexes, digested unbound RNA (schematic in Fig. 1a), and prepared cDNA libraries containing small (∼30 nt AGO2-miRNA) and large RNAs (∼70 nt AGO2-target mRNA) (Fig. 1b). To identify conserved AGO2 binding sites, we performed high throughput sequencing of three libraries for each cell type and selected sequence reads shared in all six samples. We aligned these AGO2 binding sites to human miRNA and genome databases, and identified 30-70 nt interval (peaks) significantly enriched above background (*p*-value < 0.05, Fig. 1c and methods). This analysis uncovered 316 AGO2-binding peaks within the 3’UTRs of 127 human genes. These peaks were preferentially located right after the stop codon or right before the polyadenylation site (Fig. 1d and e, Supplementary Table 1), consistent with the enrichment of regulatory miRNA binding sites that destabilize mRNAs ^24^. Importantly, the human AGO2-binding peaks within these 30-70 nt sequences showed high conservation across hundreds of species (Fig. 1e), suggesting functional significance.

**Figure 1.**
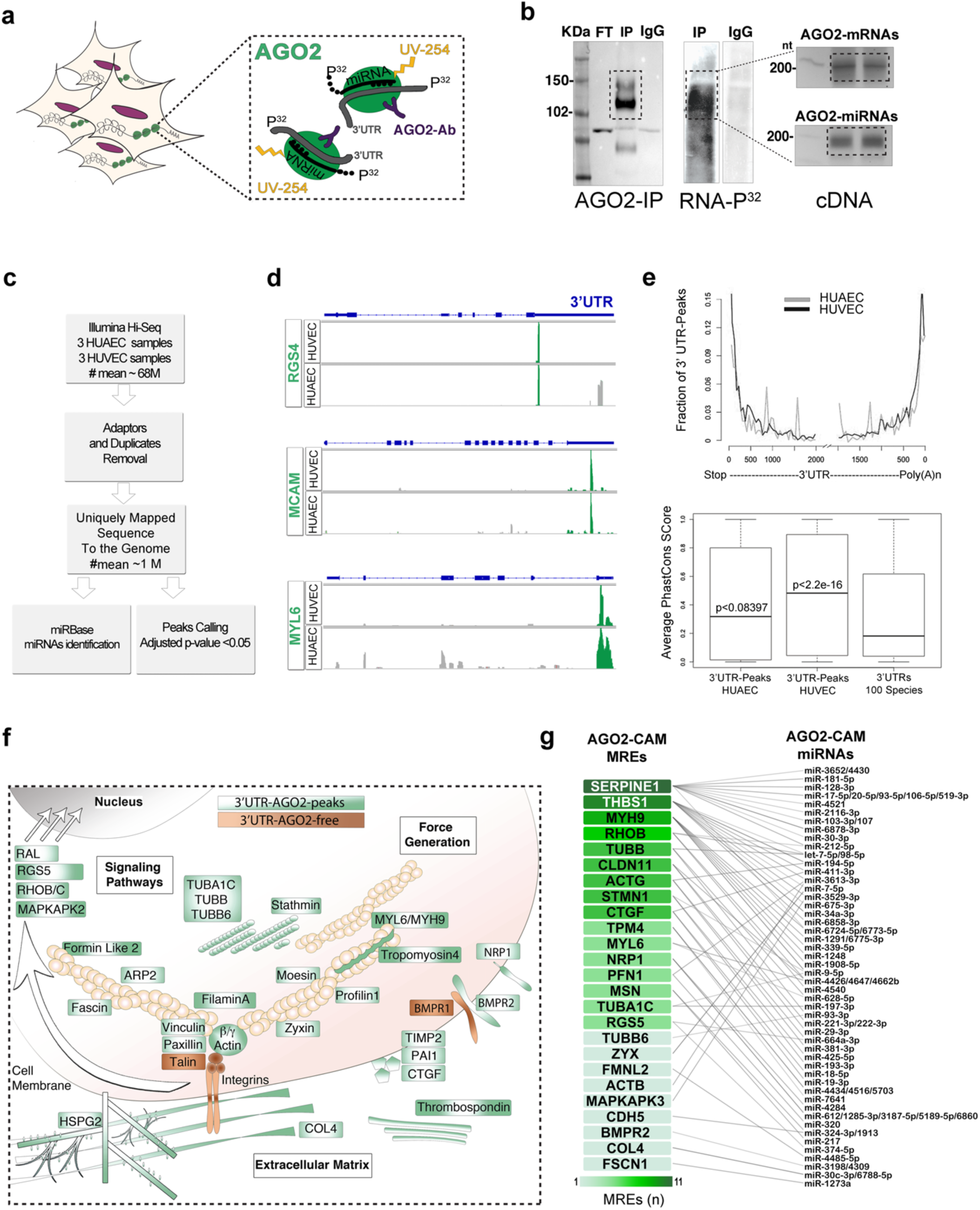
miRNA-cytoskeleton-adhesion-matrix (CAM) interactions in endothelial cells. **(a)**Schematic of AGO2-HITS-CLIP approach. miRNA-RNA complexes were cross-linked to AGO2 (green) via 254nm-UV light. RNAse treatment removed unbound RNA. P32-radiolabeling allowed isolation of radioactive RNA from AGO2-immunoprecipitated complexes (purple).**(b)** SDS-PAGE of RNA-AGO2 or control IPs with nonspecific IgG were transferred to nitrocellulose and exposed to X-Ray film to reveal P32-RNAs bound to AGO2 (dotted square). P32 RNAs were isolated (dotted line) from nitrocellulose membranes and cDNAs generated with specific primers containing an Illumina barcode for high throughput sequencing (Methods). AGO2-mRNA complexes appear as ∼200 nt bands (70 nt AGO-mRNA+ 120 nt Illumina primer), while AGO2-miRNAs are ∼150 nt (30 nt miRNA+120 nt Illumina primer). DNA MW markers are indicated in the left lane of the gel.**(c)** Computational pipeline to identify reads from miRNAs and mRNAs in AGO2 IPs. Three replicates for each cell type (HUAEC and HUVEC) were processed as described in**a** and**b.** 68 million (M) total reads were analyzed. Upon removal of Illumina adapters and duplicates from PCR-based library preparation, reads were mapped to the human genome (UCSC hg19), resulting in 1M unique reads for each cell type. Reads were mapped to miRBase to identify miRNAs and processed using Piranha software to identify significant AGO2-binding sites (peaks) (methods).**(d)** Integrative Genomics Viewer (IGV) display of HITS-CLIP reads. Reads accumulated within 30 to 70 nt intervals (AGO2 peak) within the 3’UTR region of the representative genes. Both HUAEC and HUVEC share the most significant peaks (green) while reads mapping outside the 3’UTR gene region (gray) accumulated below backgrounds (supplementary table).**(e)** Top chart represents positional enrichment of AGO2 peaks within the human 3’UTR for HUAEC (light gray) and HUVEC (black). Lines indicate the nt positional distribution of peak sequences within meta-gene analyzed 3’UTRs (methods). We observed a positional bias for AGO2 peaks at the 5’ and 3’ ends of 3’UTRs, which are typically associated with increased miRNA functionality. Bottom chart shows difference in conservation scores across samples scoring using PhastCons (Wilcoxon Rank Sum Test). AGO2 peaks in HUAEC and HUVEC and binned human 3’UTRs were compared with binned 3’UTRs of 100 species. Conservation score is represented as a boxplot. AGO2 peak sequences showed a higher conservation score compared to randomly binned human 3’UTRs.**(f)** The cytoskeleton-adhesion-matrix (CAM) AGO2-regulome. AGO2-mRNA targets identified in**a-c** highlighted in green. Integrins, Talin1 and BMPR1 proteins (boxed in brown) are part of CAM’s GO term but were not detected in AGO2-HITS-CLIP complexes. Arrows point to downstream regulators of CAM proteins targeted by AGO2. CAM proteins and their regulators were identified by database searches (Supplementary Fig. 1a) and manually curated for accuracy.**(g)** Interactome showing 25 of 73 AGO2-CAM genes in which a complementary MRE (7-8 nt) was identified using Target Scan v.7.0 prediction software and miRNAs were identified from AGO2-HITS-CLIP reads using miRbase (methods). Color-coded boxes indicate the number of MREs identified in each of the selected CAM-gene 3’UTRs. Interactions indicate interaction between MRE and miRNA family members with similar SEEDs. The mRNA-miRNA network shows high complexity in which numerous miRNAs bind one or more CAM-3’UTRs, while most CAM genes are targeted by more than one miRNA.

Gene ontology (GO) analysis of AGO2-bound transcripts revealed that 73 of the 127 target mRNAs encode actin- and microtubule-associated proteins, focal adhesion proteins, ECM proteins and functionally related regulatory proteins (Fig. 1f, Supplementary Fig. 1a). We termed this group the cytoskeleton-adhesion-matrix (CAM) genes. The dramatic enrichment of CAM transcripts in the AGO2 complex is not accounted for by their abundance; indeed, the most transcriptionally active genes in cultured ECs pertained to cell division (Supplementary Fig. 1b), which were under-represented in the identified AGO2-binding transcripts. No significant GO terms were associated with the remaining genes identified from the AGO2-HITS-CLIP.

We then searched for specific MRE sequences in AGO2-peaks localized in the 3’UTRs of the CAM transcripts. We identified 122 miRNA families from AGO2-HITS-CLIP (Supplementary Table 2) that recognize one or more AGO2-CAM MREs (Fig. 1g, Supplementary Table 3). Cytoscape software revealed a highly interconnected network of miRNAs binding to CAM transcripts (Fig. 1g). Altogether, these data reveal pervasive miRNA-mediated post-transcriptional regulation of multiple CAM genes in ECs.

CAM proteins are highly conserved and play crucial roles in virtually every cell type, as determinants of ECM organization and tissue stiffness ^25^. This important function led us to hypothesize that the CAM mRNA-miRNA regulome is mechanosensitive. To test this, we plated ECs on substrates of varying stiffness and used a Sensor-Seq strategy^26^ to assess post-transcriptional regulation mediated by 97 selected MREs within 51 different CAM 3’UTRs (Supplementary Table 4). For this purpose, we created an “MRE-sensor library”. Each AGO2-3’UTR-peak containing at least one MRE was cloned downstream of an mCherry reporter in a bidirectional lentiviral vector^27^ that co-expressed a GFP transcript lacking a 3’UTR (schematic in Fig. 2a). miRNAs that target the MRE thus reduce mCherry levels and decrease the mCherry/GFP ratio. ECs infected with this sensor library at low levels (to avoid multiply infected cells) were seeded for 48 hours on substrates with rigidity of 3 kPa (kilopascal) or 30 kPa, which approximate “soft” and “rigid” tissues^28^, respectively (Fig. 2a, Supplementary Fig. 2a). miRNA activity on the MRE sensors was compared with the steady state level of CAM proteins, CAM RNAs and miRNAs expression in the same cellular settings. Thus, proteomics, RNA and miRNA sequencing were assessed in parallel (Fig. 2a).

**Figure 2.**
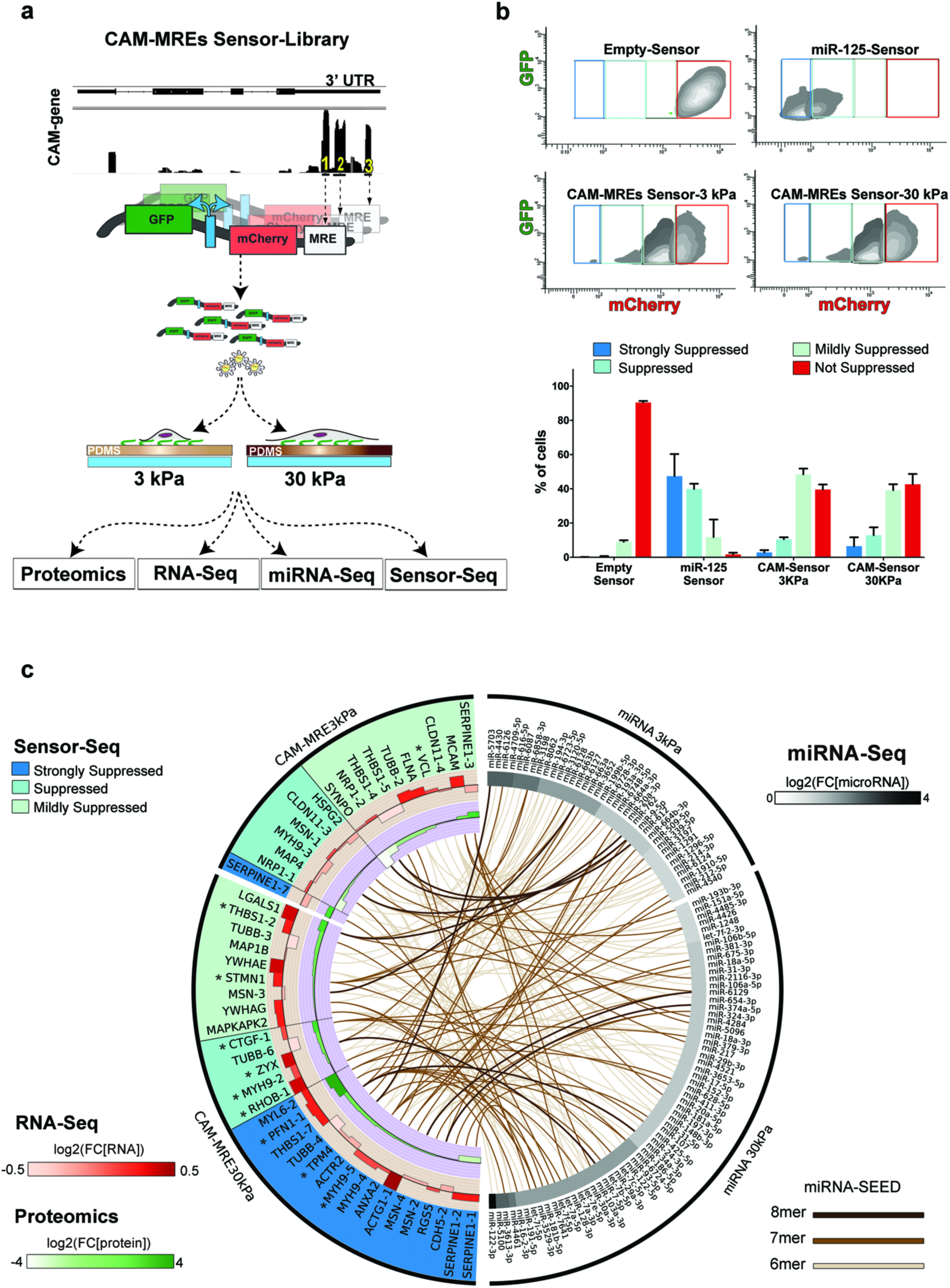
AGO2-CAM-MREs are actively repressed. **(a)**Schematic for Sensor-Seq assay of AGO2-CAM-MREs and miRNA activity. The Sensor-Seq lentiviral library (see Methods) consisted of the mCherry sequence with a 3’ noncoding sequence containing 97 CAM-MREs, plus GFP lacking any MREs as an internal control. Numbered MREs indicate different MRE positions in each clone from one CAM 3’UTR, alternatively one MRE was cloned per CAM 3’UTR (Supplementary Table 4). Titration of virus infection was performed to allow single vector copy integration and expression of physiological amounts of each GFP/mCherry transcript per cell. HUVECs were seeded on 3 and 30 kPa PDMS substrates for 48h and analyzed as described.**(b)** Intensity of mCherry and GFP in HUVECs infected with the lentiviral Sensor-Seq library; a negative control vector with no MRE, and a positive control vector with 3 perfect MREs for miR-125, which is abundant in HUVECs^74^, were included. Density plots, show relative intensity of cells distribution using contour lines. Each contour line represents 15% of probability (from higher-lighter grey- to lower-darker grey-) to contain cells in each bin over total cells (10,000 cells). Based on mCherry/GFP expression in controls, Sensor-seq library infected HUVECs at 3 kPa and 30 kPa were sorted into 4 bins as indicated. Cells in each bin were isolated and genotyped using specific Illumina primer for sequencing (Methods). The bar graph shows the percentage of cells from 4 experiments **(c)** CIRCOS^75^ graphical representation of CAM miRNA-MRE interactions. The right half of the plot shows the ECs miRNA with putative SEED matching to CAM MREs sensors differential gene expression (DEG) between 3 kPa and 30 kPa, divided in two groups: expressed at 3 kPa compared to 30 kPa (black line, top right) and expressed at 30 kPa compared to 3 kPa (black line, bottom right). The left half of the graph shows CAM-MRE Sensors most suppressed at 30 kPa compared to 3 kPa (black line, bottom left) and vice versa (black line, top left). Color-coded boxes indicate the categorized bins in **(b)** at which cells were isolated and genotyped for a specific CAM-Sensor MRE. Lines indicate match between individual miRNA (SEED) and CAM MRE in each condition. Color code indicates the level of complementarity between miRNA SEED and MRE nucleotides. The internal circles show the respective CAM-RNAs (red) and proteins (green) log2 fold change at 3 kPa compared to 30 kPa (top right) and 30 kPa compared to 3 kPa (bottom right).

To assess CAM-MREs sensor reporters, ECs were separated by fluorescence-based sorting into bins according to the mCherry/GFP ratio, using an empty-Sensor as a negative control (not suppressed) and a miR-125-Sensor as a positive control (strongly suppressed). Thus, bins were defined as strongly suppressed, suppressed, mildly suppressed, and not suppressed relative to these internal standards (Fig. 2b). Wild-type ECs infected with our CAM-MREs Sensor library showed a broad distribution between the suppressed and not suppressed bins, on both soft and stiff substrates (Fig. 2b). Importantly, CRISPR/Cas9-mediated disruption of AGO2 diminished the miRNA levels in CAM-MREs Sensor ECs (Supplementary Fig. 2b and c), and significantly increased the population of ‘not suppressed’ cells (Supplementary Fig. 2d). Thus, miRNAs are required for post-transcriptional inhibition of CAM-MRE Sensors.

Sensor vectors from sorted cells were then isolated from each bin and barcoded using PCR primers that recognized each cloned CAM MRE and were compatible with high throughput sequencing (Sensor-Seq). Combining global miRNA profiling and MRE-reads from Sensor-Seq revealed strong correlations between suppression of CAM MRE sensors and the level of the respective matching miRNAs (Fig. 2c). Notably, both miRNA levels and CAM reporter suppression were present on soft substrate at baseline and elevated in cells on stiff substrates (Fig. 2c). Interestingly, the levels of most CAM mRNA and respective proteins were also generally higher in stiff conditions (Fig. 2c). These results suggest transcriptional co-regulation between miRNAs and CAM mRNA targets on stiff substrates. Thus, the CAM MRE-miRNA network has the characteristics of a mechanoregulatory buffer of structural protein coding genes.

To evaluate the function of this miRNA regulatory network, we first examined ECs lacking AGO2 or DROSHA, which have diminished miRNA levels (Supplementary Fig. 2 b and c and ^29^). We used immunofluorescence to detect F-actin, the focal adhesion marker paxillin, and the mechanosensitive transcription factor YAP^30^, and also assayed for traction stresses using elastic substrates with embedded beads^31^. Relative to control cells, AGO2 mutant cells showed increased actin stress fibers, focal adhesions, Yap nuclear localization and traction stress on both 3 or 30 kPa substrates (Fig. 3a), as well as on polyacrylamide substrates over a wider range of stiffnesses (Supplementary Fig. 3a and b). Consistent with these observations, immunofluorescence analysis revealed that cell spreading and YAP nuclear activation were inversely correlated with AGO2 levels (Supplementary Fig. 3c). Furthermore, proteomic analysis of AGO2 mutant cells showed increased levels of several CAM proteins, reminiscent of the increased CAM levels in ECs plated on 30 kPa versus 3kPa substrates (Fig. 3b and Supplementary table 5). Importantly, human dermal fibroblasts deficient in AGO2 protein had similar hyper-adhesive and contractile behaviors, as did ECs lacking DROSHA (Supplementary Fig. 3d and e). Together these data suggest that the loss of miRNA-mediated suppression of mRNAs increases CAM protein levels and enhances cell contractility and adhesion in different cell types.

**Figure 3.**
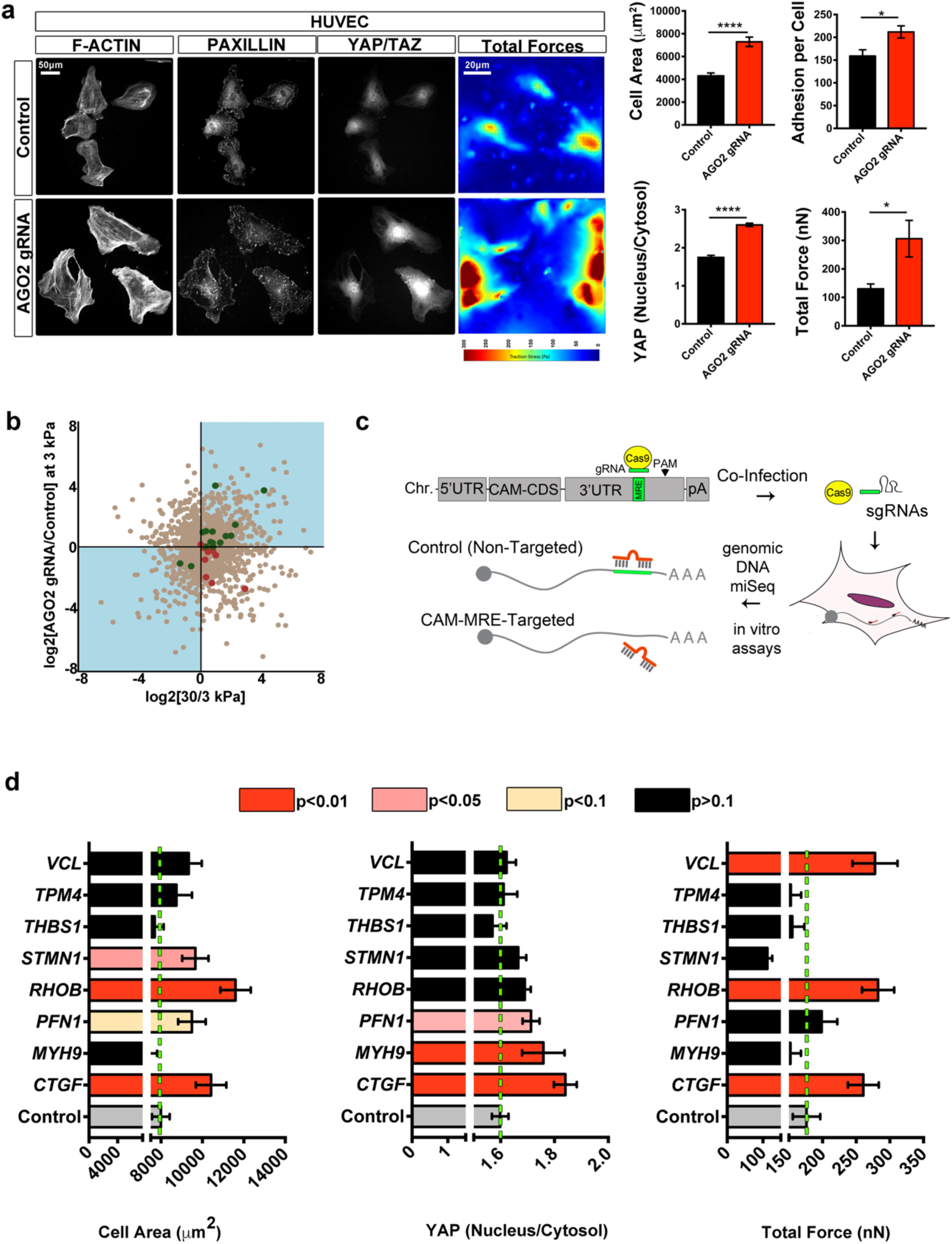
miRNA interactions limit cell spreading, YAP signaling and contractility. Representative immunofluorescence images and traction force maps of HUVECs (a) after infection with AGO2 or non-targeting control pLentiCRISPR virus. Cells on fibronectin-coated 3kPa PDMS gels were stained for F-Actin (phalloidin), focal adhesions (paxillin), and YAP/TAZ. Quantification shows cell area (based on phalloidin staining) (n=281-396 cells/group), number of paxillin adhesions per cell (n=49-51 cells/group), and nuclear to cytoplasmic ratio of YAP/TAZ (n= 43-54 cells / group). Single cell maps of average traction stress and quantification of total force per cell (n=19 cells / group, bars indicate standard error, * p<0.05, **** p<0.0001,). **(b)** Scatter plot representing difference in proteins expression between HUVEC seeded on 30 vs. 3 kPa (x axis) or between HUVECs infected with AGO2 gRNA vs. control gRNA (y axis). Green and red identify CAM proteins with coherent or incoherent differential expression, respectively (Supplementary Table 6). **(c)** Experimental strategy to mutate individual MREs in CAM gene 3UTR’s to block miRNA binding (See methods and Supplementary Figs 4 and 5). **(d)** Quantification of cell spreading (n=68-99 cells/group), YAP nuclear translocation (n=68-99 cells/group) and traction stress (n=13-21 cells/group) in HUVECs on 3 kPa PDMS gels for 48h after mutation of the indicated MREs (bars indicate standard error).

To further validate the function of the miRNA-CAM mRNA network, we disrupted individual CAM-miRNA interactions. We chose nine of the mechanosensitive CAM MREs (stars in Fig. 2c, and Supplementary table 6) in which the MRE was within 20 nt of a protospacer sequence (PAM) and thus targetable by a guideRNA (gRNA) and Cas9. Genome-wide analyses of CAM MREs in ECs treated with gRNA/CRIPSR/Cas9 revealed insertions and deletions within the desired MRE region (Fig. 3c and Supplementary Figs. 4 and 5). We found that mutation of individual CAM MREs de-repressed CAM protein levels, as predicted for impaired miRNA-mediated suppression^32^, and increased, to varying extents, cell area, YAP nuclear localization and/or traction stresses (Fig. 3d, Supplementary Fig. 6a).

While multiple genes clearly contributed to each effect, the gene whose MRE mutation gave the most consistent effects across multiple assays was Connective Tissue Growth Factor *(CTGF).* CTGF is a matrix protein that modulates the interaction of cells with the ECM^33^, suggesting that it is a component of a protein-based regulatory network and likely functions via receptor-mediated signaling to control these functions. Blocking CTGF miRNA-repression in ECs via a target protector RNA oligonucleotide or MRE mutation had similar effects (Fig. 3d, Supplementary Fig. 6 a and b), providing independent support. Notably, no single MRE mutation reproduced the strong phenotype observed after AGO2 downregulation, suggesting that a network of miRNA-CAM mRNA interactions mediates mechanical homeostasis in cells.

As a first approach to determine if the miRNA-mediated network functions at the tissue level, we examined primary mouse dermal fibroblasts grown in a 3-dimensional (3D) matrix. Cells suspended in attached fibrin gels contract and replace the fibrin with their own matrix over about 5 days (Fig. 4a), providing a 3D model of cell behavior. Transduction of these cells at passage 0 with a CRISPR/Cas9/gRNA virus targeting *Ago2* reduced Ago2 protein levels by ∼50-60% (Supplementary Fig. 7a). Ago2-depleted fibroblasts grown in 3D matrix generated tissue constructs with reduced diameters but no significant change in cell numbers (Fig. 4b). Immunostaining transverse sections of these constructs confirmed the decreased diameter, based on staining with the cytoskeleton protein Vimentin (Fig. 4c). We observed elevated staining for phosphorylated myosin light chain, demonstrating that reduced Ago2 levels led to increased myosin activation and contractility^34^ (Fig. 4c). These data suggest that reducing miRNA-dependent regulation stimulates contractility in fibroblasts in a 3D setting.

**Figure 4.**
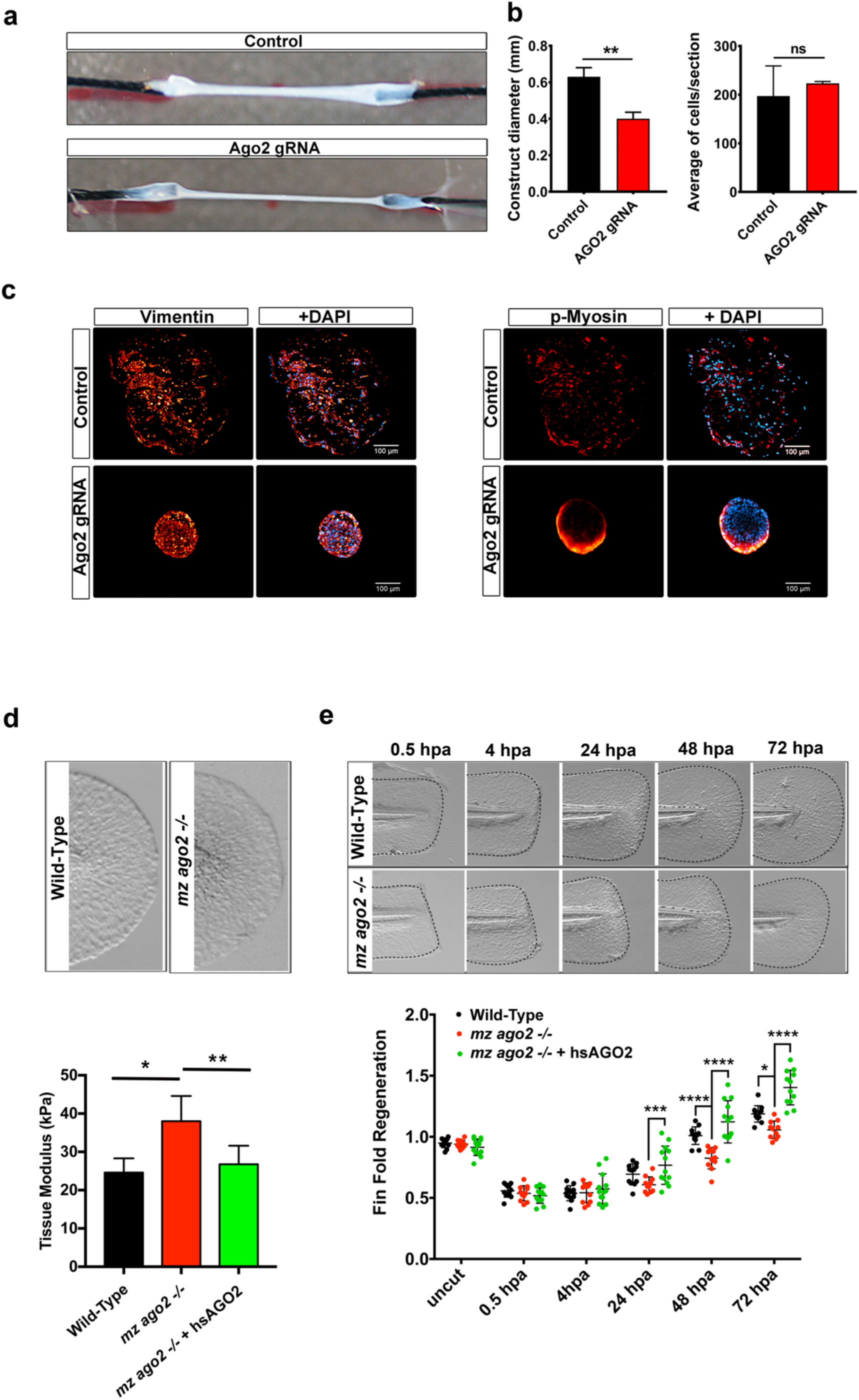
AGO2 activity limits tissue contractility and stiffness. (a)Representative 3D matrix constructs with control or Ago2-mutated mouse dermal fibroblasts. **(b)** quantification of cell number and construct diameter within transverse sections (n=8, bars indicated standard error, ** p<0.01, ns= non significant). **(c)** Transverse sections of control and Ago2 depleted matrix constructs stained for Vimentin or pMyosin. **(d)** Bright filed images of fin fold tissues in the indicated genotypes. Elastic modulus of 48 hpf zebrafish fin fold surfaces of Wild-Type, *mz ago2 -/-* and *mz ago2 -/-* rescued with 200 pg of *in vitro* transcribed human Wild-Type AGO2 mRNA (hs *AGO2 mRNA).* Embryos were harvested and adhered to a soft surface of PDMS in egg water. Elastic moduli were measured and AFM using NanoScope Analysis 1.5 software to fit force-deflection curves using the Sneadon model. At least two fin fold regions within each of 10-11 embryos were tested for each genotype (n=27-49 measurements per region; bars indicate standard deviation, *p<0.05). **(e)** Bright field images of zebrafish fin folds at the indicated stages and genotypes (head is to the left). Dotted black line outlines the edge of the fin fold. Fin fold regeneration was assessed from the distance between the wound edge and the embryo body. One cell stage Wild-Type, *mz ago2-/-,* and *mz ago2-/-* embryos rescued with 200 pg of *hs AGO2 mRNA.* Values were normalized for the size of the fin-fold prior to injury (n=14, bars = standard deviation, *p<0.05, ** p<0.01; ***p<0.001; ****p<0.0001).

We next tested whether miRNAs regulate mechanical homeostasis *in vivo* using the zebrafish fin fold regeneration model^35^. The fin fold is a non-vascularized appendage comprised of a few layers of epidermis and fibroblast-like cells^36^. Wounding triggers a healing response mediated by a conserved and rapid matrix remodeling- and actomyosin-based process that involves formation of a provisional matrix, inflammatory cell invasion, cell migration, proliferation and resolution^37^.

To investigate miRNA-dependent regulation of mechanical homeostasis in zebrafish, we first examined embryos that carry a maternal zygotic homozygous mutation in *ago2 (mz ago2* -/-)^38^, which show reduced levels of Ago2 and of miRNAs (Supplementary Fig. 7b). To evaluate miRNA activity in the fin fold of *mz ago2* -/-embryos, we co-injected an miRNA-sensitive GFP mRNA, containing three perfect miR-24 MREs within the 3’UTR^39^, with an miRNA-insensitive mCherry control mRNA. As expected, *mz ago2* mutants showed elevated levels of GFP, but not mCherry, when compared to wild-type (WT) embryos, confirming reduced miRNA-mediated suppression (Supplementary Fig. 7c). We then quantified tissue stiffness using atomic force microscopy (AFM)-based nanoindentation on the central region of the fin fold. The appearance of this tissue was indistinguishable between genotypes (Fig. 4d), ruling out obvious developmental defects. However, the elastic modulus was ∼30% higher in *mz ago2* -/-than WT embryos, indicating increased mechanical rigidity (Fig. 4d). Importantly, normal tissue stiffness was restored upon injection of *in vitro* transcribed mRNA encoding human AGO2 *(hsAGO2 mRNA),* demonstrating that the stiffness of this tissue is dependent upon the level of Ago2 (Fig.4d). Following amputation, *mz ago2* mutants exhibited slower repair than WT embryos and *mz ago2 -/-* embryos expressing *hsAGO2 mRNA* (Fig. 4e). WT and *mz ago2* -/-wounds did not display differences in cell cycle progression, detected by Proliferating Cell Nuclear Antigen (PCNA) staining^40^, or in apoptosis, detected by TUNEL assay (Supplementary Figs. 8 a and b and 9a). These results support that miRNA-dependent suppression is required mainly to restrain tissue stiffness and contributes to tissue healing *in vivo.*

Wounding triggers increased contractility and matrix rigidity as a rapid, first response. According to our notion of mechanical homeostasis, these changes should activate negative feedback mechanisms that restore mechanical equilibrium^3^. We therefore examined matrix, actomyosin activation and the mechanosensitive translocation of Yap^30^ before and after wounding the zebrafish fin fold in WT vs. *mz ago2* mutant. As expected^37^, WT embryos showed increased staining for pMyosin, Ctgfa and Fibronectin^34,37^ in the wound area between 0.5 and 2 hours post amputation (hpa), (Fig. 5a and b, Supplementary Figs. 8 c and 9a). In comparison, *mz ago2* -/- wounded fins showed strikingly elevated and persistent pMyosin staining at both 0.5 and 2 hpa, and higher Ctgfa and Fibronectin at 2 hpa (Fig. 5a and b, Supplementary Figs.8c and 9a). Consistent with the increase in tissue stiffness (Fig.4d), *mz ago2* -/- showed higher basal Yap nuclear localization compared to WT embryos that further increased at 0.5 hpa and persisted at 2 hpa after wounding (Fig.5c and Supplementary Fig. 9b). Thus, loss of miRNA-mediated suppression leads to an exaggerated mechanical response and impaired mechanical resolution during wound healing.

**Figure 5.**
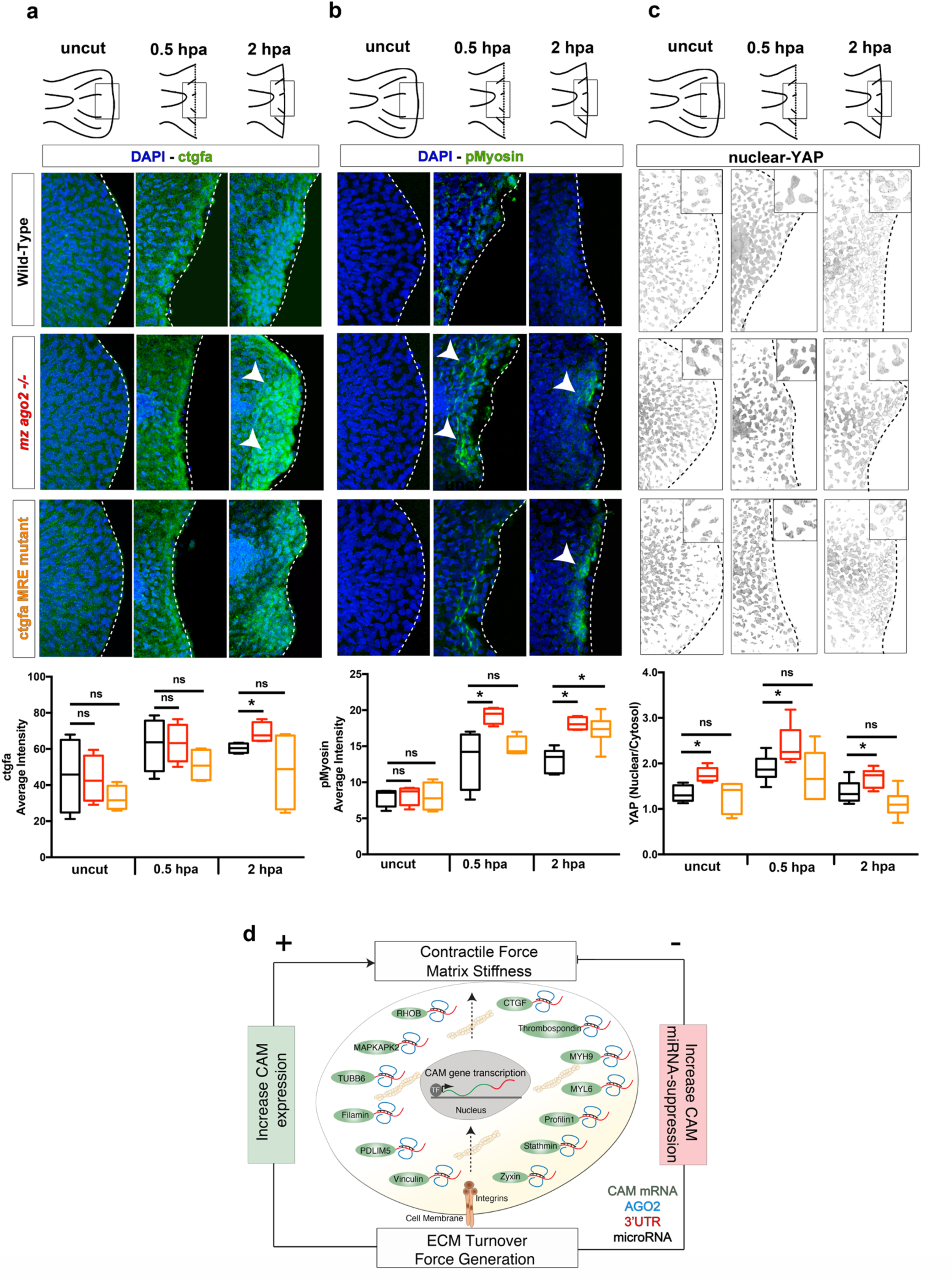
Wound healing in mz *ago2* and ctgfa MRE fin fold mutants. (a-c)Top, schematics representing the time course of fin fold regeneration. Boxes identify the region of interest reported in the images below. Bottom confocal images of whole mount fin folds within the boxed region from the cartoons at the top, at the indicated stages. White and black dotted lines indicate the edge of the fin fold. White arrows point to staining for the indicated markers. Graphs show box plot of the respective measurements (methods and Supplementary Fig. 9a). Intensity profiles for multiple embryos were combined (n= 6 embryos for each genotype, *p<0.05, n.s.= non significant). For Yap the protein nuclear localization was represented. The nucleus/cytosol ration was obtained using the DAPI channel to generate a binary mask and subtract nuclear YAP-GFP intensity from the total YAP-GFP detected (methods and Supplementary Figure 9b). **(d)** The cartoon represents a model for miRNA-post-transcriptional regulation of structural proteins function in mechanical tissue homeostasis. Increases in matrix stiffness and the resulting cell contractility increase integrin and actomyosin-dependent CAM signaling, which upregulates miRNAs that suppress CAM transcripts, thus restoring normal tissue mechanics.

To correlate these effects to regulation of individual CAM genes, we generated zebrafish embryos carrying mutations in the two 3’UTR MREs of the *ctgfa* gene (Supplementary Fig. 8 d and e). These MREs are conserved in the human *CTGF* 3’ UTR, and their mutation had the largest effect *in vitro* (Fig. 3d, Supplementary Figs. 4 and 6). Accordingly, a GFP sensor mRNA bearing a *ctga* 3’UTR fragment showed reduced expression in WT relative to *mz ago2 -/-* embryos (Supplementary Fig. 8 d), which required the MRE sites (Supplementary Fig. 8 d). These results support miRNA-dependent inhibition of *ctgfa* via the MREs in zebrafish. We found that zebrafish with mutated *ctgfa* MREs showed persistent p-myosin activation compared to WT by 2 hpa (Fig. 5b), consistent with the induction of Ctgfa at 2 hpa in the *mz ago2* mutant (Fig. 5a). However, no other differences were detected in the *ctgfa* MRE mutant embryos (Fig. 5a-c, and Supplementary Figs.8 a-c and 9). These results support that post-transcriptional regulation of *ctgfa* contributes to specific Ago2-mediated mechanical effects within the miRNA-CAM mRNA network.

## Discussion

We report that an unbiased analysis of miRNAs and their target genes in endothelial cells, together with functional assays in several biological systems, reveal the existence of a mechanosensitive miRNA-based program that counteracts cell adhesion, cytoskeletal, contractile and matrix protein expression. This system is highly conserved, functioning in several cell types, across multiple species, and with evidence of high evolutionary conservation. Importantly, most of the protein-coding genes for synthesis and assembly of stiff ECM are targeted by miRNAs on stiff substrates. Thus, a “buffer” is generated, in which increased matrix stiffness not only upregulates cytoskeleton-adhesion-matrix gene transcription, but also upregulates miRNA-mediated suppression of cytoskeleton-adhesion-matrix transcripts. This miRNA-regulome has the molecular and functional characteristics of a homeostatic mechanism in which changes in cell contraction and matrix are counteracted to maintain normal tissue stiffness (Fig. 5c).

A network-mediated mechanism for stiffness homeostasis, rather than regulation of one or a few CAM genes, would be expected to increase the robustness of the system. Multiple miRNAs can regulate a large cohort of CAM genes via different MREs, while different cell types can do so by controlling expression and processing of tissue specific mature miRNAs^41-43^. We speculate, however, that these miRNA networks are likely to be sub-elements within a larger and more robust network of negative and positive circuits, connected by multiple nodes, that mediate tissue homeostasis over the multiple decades of human life^44^. Such nodes could develop within a hierarchy of epigenetic factors in which, for example, the activation of YAP/TAZ and its direct target gene *CTGF,* may be one of the upstream components.

A role for miRNAs in tissue mechanical homeostasis is supported by the widespread de-regulation of miRNAs in lung, renal, cardiac and liver fibrosis, including miRNAs that target ECM proteins^45-47^. Idiopathic lung fibrosis is also linked to reduced levels of miRNAs that target ECM, cytoskeletal and TGFß pathways genes ^48-50^. All of these studies reported reduced levels of miR-29 species, in contrast to our finding that in normal cells, miR-29 species are increased on stiff substrates. These results are consistent with the notion that fibrotic disease involves disruption of normal stiffness miRNA-dependent homeostasis^51^.

miRNA-dependent post-transcriptional regulation of structural proteins provides a concrete molecular mechanism that can explain how healthy tissues sustain optimal mechanical properties. These findings are therefore an important step toward understanding the initial pathological alterations resulting in fibrotic and related diseases. Characterizing the stiffness-dependent RNA metabolism of cytoskeleton and matrix transcripts, their possible regulation under other physical forces, and elucidating the complete regulatory network that mediates long-term mechanical robustness are the essential tasks for future studies.

## Methods

### Cell Culture

Human umbilical vein endothelial cells (HUVECs) and human umbilical artery endothelial cells (HUAECs) were purchased from Cell Applications Inc. (Cat # 200-05n and Cat #202-05n). Endothelial cells were cultured on tissue culture dishes coated with 0.2% w/v gelatin (10 min at room temperature in PBS, Sigma) in endothelial cell growth medium (EGM Bullet Kit, LONZA). For HITS-CLIP assays, cells were used at P3 (split 1:3 and 1:5) before UV crosslinking. For other assays, cells were split 1:3 twice per week and used until passage 5. Human dermal fibroblasts (HDFs) from normal donors were purchased from ATCC (Cat #PCS-201-010, Lot# 63014910) and cultured on 0.2% w/v gelatin-coated dishes in fibroblast growth medium (Fibroblast Growth Kit-Low Serum, ATCC, PCS-201-041). HDFs were split 1:10 twice per week and used until passage 6.

### Primary fibroblasts

Primary dermal fibroblasts for 3D fibrin gel assays were obtained from 5 to 8 week old C57BL/6 mice (Envigo, UK). All procedures were in accordance with UK Home Office regulation and UK animals (Scientific Procedures) Act of 1986 for the care and the use of animals. Mice were sacrificed by a schedule 1 procedure by trained personnel. Mouse hair was removed with a hair clipper and skin dissected in Hank’s buffer supplemented with antibiotic and antimycotic solution (Sigma). Fat and excess connective tissues were removed, the dermis was minced with a scalpel and digested in buffer containing 0.25% trypsin without EDTA (Gibco), collagenase IV (4mg/mL (Worthington) and calcium chloride (0.3mg/mL, Sigma) for 3 hours at 37°C with frequent agitation during the last hour. After mechanical dissociation, cells were passed through a cell strainer (100 μm, Fischer Scientific) to remove debris and hairs. Cells were centrifuged at 1800 rpm for 5 minutes, resuspended in DMEM supplemented with 10% Fetal Bovine Serum (Sigma), Penicillin (100U/mL), Streptomycin (100 ug/mL) (Gibco) and 1% L-glutamine, and seeded in 75cm^2^ tissue culture flasks. Medium was changed at 3 hours and subsequently changed once a day.

### AGO2-HITS-CLIP

HITS-CLIP experiment was performed as previously described ^22^. Three independent replicas were analyzed for each cell line (HUVEC and HUAEC). For each replica, five 10 cm dishes of sub-confluent endothelial cells (ECs) in EGM Bullet Kit supplemented media (LONZA) were UV crosslinked two times with 400mJ/cm^2^ in Stratalinker (model 2400, Stratagene), lysed and treated with DNase (1:1000 Promega RQ1 DNase) and RNase T1 (1:100, Thermo Fisher). Cell lysates and Protein A Dynabeads (Invitrogen) complexed with Ab-panAGO-2A8 (MABE56 Millipore) were incubated at 4°C for 4 hours. Beads were subsequently washed and ligated with 3’ -P32 radiolabeled linker (RL3: 5’-PGUGUCAGUCACUUCCAGCGG-3’). SDS-PAGE was performed using NuPage 4-12% Bis-Tris Gel (NP0321 Invitrogen), and proteins were transferred onto Pure Nitrocellulose membrane (BioTrace) using NuPAGE transfer buffer according to manufacturer’s instructions. High performance autoradiography film was exposed overnight at −80°C. The bands corresponding to AGO2-miRNAs (∼110 kDa), and AGO2: RNA (∼130 kDa) were cut and treated with proteinase K (Roche) to degrade proteins. RNAs were extracted and purified via phenol-chloroform, then a 5’-linker oligonucleotide (RL5: 5’-AGGGAGGACGAUGCGG-3’) was ligated to the ends. cDNA libraries were generated using DNA oligos complementarity to RL3 and SuperScriptIII reverse transcriptase (Invitrogen) according to manufacturer’s instructions. Products were then PCR amplified using specific primers (DP5: AGGGAGGACGATGCG, DP3: GCCGCTGGAAGTGACTGACAC) and purified via agarose PAGE 1% using a Gel extraction kit (Qiagen). A second round of PCR was performed, using custom Illumina Hi-Seq primers with three different barcodes to multiplex the libraries (DSFP5 5’ to 3’ sequence: AATGATACGGCGACCACCGACTATGG, DSFP3-Index1: CAAGCAGAAGACGGCTATCGAGATTGGTCAGTGACTGGAGTTCACCGCTGGAAGTGACTGACAC, DSFP3 5’ to 3’ sequence -Index4:

CAAGCAGAAGACGGCTATCGAGATCGTGATGTGACTGGAGTTCACCGCTGGAAGTGACTGACAC

DSFP3 5’ to 3’ sequence -Index8

CAAGCAGAAGACGGCTATCGAGATTCAAGTGTGACTGGAGTTCACCGCTGGAAGTGACTGACAC). Products were PAGE purified using the Gel extraction kit, and libraries were analyzed by the YCGA Sequencing facility using a customized Illumina primer (SSP1 5’ to 3’ sequence: CTATGGATACTTAGTCAGGGAGGACGATGCGG).

### Data Analysis

Human AGO2 peaks were called using Piranha peak caller (version 1.2.1)^52^. Prior to aligning sequencing reads, the raw data were analyzed for quality steps to reduce artifacts: adapters were removed from raw reads, filtered according to quality scores and exact sequence duplicates were collapsed. Remaining reads were aligned using STAR RNA-seq aligner (version 2.4.1a)^53^ using UCSC hg19 reference human genome. A minimum of 10 bases matched was enforced, only unique reads were used, and a maximum of 3 mismatches were allowed. Replicates were merged using Samtools (version 1.2) ^54^ and the aligned reads were analyzed with Piranha using a bin size of 30 bp. All identified peaks with p-value less than 0.05 were mapped to Gencode version 22 annotation.

Conservation between artery and vein samples was calculated for each identified peak using PhastCons 100 conservation scores. Using Piranha peaks with a 30bp bin setting, a Wilcoxon rank sum test was performed to compare the difference in distribution of conservation score across samples.

For microRNA identification, reads were aligned using Novoalign (Novocraft, http://www.novocraft.com/products/novoalign/) against human microRNA sequences from miRBase (release 21)^55^. The miRNA expression levels were quantified as the number of reads mapped to individual miRNA genome sequence and normalized to the total number of mapped reads in miRbase per million (RPM). Endothelial microRNAs identified in AGO2-HITS-CLIP were divided into families based on 8mer SEED regions. Using TargetScan software ^56^, these microRNA SEED families were associated to the AGO2-HITS-CLIP peaks based on the putative MRE.

To test expression of CAM vs. other genes in cultured ECs, we examined previously published microarray data performed in freshly isolated versus cultured HUVEC and HUAEC (GEO ID: GSE43475) ^57^. Standard microarray analysis for differential gene expression (DEG) was performed using Bioconductor library of biostatistical packages (http://www.bioconductor.org/) and specific packages simplyaffy (http://bioinformatics.picr.man.ac.uk/simpleaffy/) and limma ^58^.

### CRISPR/Cas9 strategy to generate mutant primary cells

To mutate *AGO2* in HUVECs and HDFs, a pLentiCRISPR vector containing an AGO2 or a non-targeting guide RNA (control, which doesn’t target know mouse or human genome sequences) was used (AGO2(Fw): 5’-CACCGGGGGCCGGCTCCCGAGTACA-3’, AGO2(Rv):5’-

CACCGGCGTTACACGATGCACTTTC-3’) (NT(Fw): 5’-GCGAGGTATTCGGCTCCGCG, NT(Rv): 5’-CGCGGAGCCGAATACCTCGC-3’). Lentiviruses were generated by transfecting Lenti-X 293T cells (Clontech) (10cm dishes) with packaging vectors (2.5μg VSV; 5μg pxPAX2, Addgene) and the pLentiCRISPR DNA vector (7.5ug) using lipofectamine 2000 (Invitrogen). Virus containing supernatant was collected 36 and 60 hours post transfection. In Supplementary Fig. 2b shows a schematic of approach. Cells were infected with pLentiCRISPR virus containing AGO2 gRNAs or non-targeting gRNAs in the presence of polybrene (8 μg/ml). To generate cells with mutant MREs, similar vectors were generated to target the selected MRE sequences identified in the AGO2 HITC-CLIP experiment and confirmed via Sensor-seq. The complete lists of genes and gRNAs are reported in Supplementary Table 6. Cells were cultured for 7-10 days (up to a maximum of passage 5) prior to seeding on gels for immunostaining or traction force microscopy. Reduced AGO2 expression was confirmed by Western blot at 7 and 10 days post-infection. Cells were lysed in RIPA buffer with protease and phosphatase inhibitor cocktail (Roche) on ice. Samples were loaded onto 8% or 4-12% SDS-PAGE gels and run 2h at 130 V. Transfer was performed using Tris-Glycine/Methanol buffer on Immun-Blot PVDF membrane (Biorad). Membranes were blocked with 10% milk for 2 hours and incubated with rabbit anti-AGO2 (Cell Signaling) and mouse anti-βActin (Santa Cruz) in 2% milk overnight at 4°C. After washing, membranes were incubated with secondary antibody anti-mouse-HRP and anti-rabbit-HRP (Santa Cruz) for 1h. For blots of other proteins following MRE mutation, target protector or knockdown, membranes were blocked with 5% w/v BSA in PBS 0.1% Tween for 1 hour and incubated with primary antibody for RhoB (1:200, sc-8048, Santa Cruz), CTGF (1:1000, ab6992, Abcam), Vinculin (1:2500, V9131, Sigma-Aldrich), STMN1 (1:10000, ab52630, Abcam), DROSHA (1:5000, ab183732, Abcam), or GAPDH (1:4000, 2118, Cell Signaling) overnight at 4°C. After washing, membranes were incubated with secondary antibody anti-rabbit-HRP or anti-mouse-HRP (1:4000, 7076P2 and 7074S, Cell Signaling) for 1 hour at room temp in 5% BSA TBS-T. After washing, blots were developed with super signal west pico chemiluminescent substrate (Thermo) using a SYNGENE G-Box imager.

For single CAM MRE mutations, T7 Endonuclease I assay^59^ was first used to verify the occurrence of indels in the MRE sequence as described in manufacturer’s protocol (New England BioLabs)(data not shown). Then, single amplicons of ∼300 bp were generated using primes equidistant from the putative region of mutation. PCR amplicons were combined and sent to Yale Sequencing Facility for MiSeq 2×250 analysis. After Illumina sequencing, single amplicons were demultiplexed and single reads were used for msa (Multiple sequence alignment) against the wild-type sequence using R msa package^60^. The frequency of each mutation was calculated as total reads for each CAM gene mutation divided the sum of all the reads aligned to specific CAM gene, and plot as bar plot.

To mutate *Ago2* in mouse fibroblast, pLentiCRISPRV2 (Addgene) vectors containing Ago2 or non-targeting guide RNAs (control, as above) were used. P0 fibroblasts at 80% confluence were infected with lentivirus containing either non-targeting or Ago2 guide RNA in the presence of 4mg/mL polybrene for 16 hours. Culture medium was changed and cells were incubated for 72 hours. Infected cells were selected in medium with 0.5 ug/mL puromycin for 48 hours (this concentration efficiently kills all control cells) and then cultured for another 96 hours before use in matrix constructs. Reduced Ago2 expression in mouse fibroblasts was confirmed by Western blot of protein extracts from the matrix constructs at 5 days. Matrix constructs were washed with cold PBS and immediately immersed in liquid nitrogen. Frozen constructs were then homogenized with metallic beads in a Bullet Blender (Strom 24, Next Advance) in protein extraction buffer (1.1% Sodium dodecyl sulphate, 0.3% sodium deoxycholate, 25mM dithiothretol, in 25mM ammonium bicarbonate with Complete anti-protease and anti-phosphatase, Roche). Protein samples were loaded in a 4-12% Nu-Page pre-casted gel (ThermoScientific) for electrophoresis (200V, 50 minutes). Transfer was performed using Tris-Glycine/Methanol buffer on nitrocellulose membrane (Biorad). The membrane was then blocked for 1 hour with Odyssey PBS blocking buffer (LiCor biosciences) and incubated overnight with 2 primary antibodies against Ago2 (Cell Signaling) and beta-actin (Abcam) at 1/5000. After extensive rinsing in PBS-Tween, the membrane was incubated with 2 secondary antibodies: AlexaFluor 680 anti-mouse to detect α-actin and AlexaFluor 800 anti-rabbit to detect α-Ago2 (both from Thermoscientific, 1/15000). The membrane was scanned with an Odyssey CLX NIR scanner (Licor biosciences) and fluorescence intensity of the bands quantified with the Image Studio software (Licor biosciences).

### shRNA knockdown of DROSHA and miRNA Target Protector for CTGF

Knockdown of DROSHA was performed using Dharmacon shRNA SMARTvectors (GE Healthcare). Lentivirus was prepared in Lenti-X 293T cells as before using a non-targeting negative control shRNA (5’-CCTAAGGTTAAGTCGCCCT-3’) and two shRNAs directed at DROSHA (shDRO#1, 5’-ACCAATGCCTTGTCCTAAT-3’) (shDRO#2, 5’-GCAAAGGCATGATTGTTAC-3’). Experiments were performed with shDRO#2 since it achieved ∼95% knockdown at 5 days post infection. DROSHA knockdown was verified by immunoblot as before with DROSHA antibody (Abcam, ab183732, 1:5000 in 5% BSA).

Disruption of the miRNA-MRE interaction with the CTGF gene was performed using a miScript Target Protector (Qiagen) directed at the MRE within the human CTGF gene. The CTGF target protector (CTGF_1_TP, Cat#MTP0079186, 5’-AACTAGAAAGGTGCAAACATGTAACTTTTG-3’) or the Negative control target protector (Cat#MTP0000002) were transfected into P2 HUVECs at 20nM using Lipofectamine RNAiMax (Invitrogen) in OPTI-MEM (Gibco) with 4% FBS (Sigma). CTGF increases post transfection were verified by immunoblot as before with a CTGF antibody (Abcam, ab6992, 1:1000 in 5% BSA).

### PDMS and Polyacrylamide Substrates

Polydimethylsiloxane (PDMS) substrates were cast in the bottom of 10cm tissue culture dishes or #1.5 cover-glass bottomed 35mm Mattek dishes (for imaging studies). Soft (3kPa) gels were made using a 1:1 ratio (by weight) of PDMS component A and B (CY 52-276 A and B, Dow Corning), degassed for 30 minutes in a vacuum desiccator, and cured for 24 hours at room temperature. Stiff (30kPa) gels were made using a 40:1 ratio (by weight) of Sylgard 184 components B and C (SYLGARD 184, Dow Corning), degassed for 30 minutes and cured for 3 hours at 70°C. Prior to seeding, gels were washed with PBS, sterilized with UV for 20 minutes, and coated with bovine plasma fibronectin (10μg/ml in PBS) overnight at 4°C.

Polyacrylamide gels were prepared using a protocol modified from previously published methods in ref ^61^. Briefly, 30-mm glass bottom dishes were activated with glacial acetic acid, 3-(trimethoxysilyl) propyl methacrylate, and 96% ethanol solution (1:1:14 ratio, respectively) for 10 minutes in room temperature. For fibronectin protein conjugation (1mg/ml) on the polyacrylamide gel, acrylic acid N-hydroxysuccinimide ester was partially mixed as a substitute of acrylamide. Each stiffness was prepared with the ratio in the table below which was previously reported by ref ^61^.

| Stiffness (Pa) | 40% aa (ul) | 2% bis-aa (ul) | 60% NHS acrylate (ul) | Citric buffer (pH 4) (ul) | APS (ul) | TEMED (ul) |
| --- | --- | --- | --- | --- | --- | --- |
| 490 | 45 | 7.5 | 33 | 394 | 20 | 1 |
| 1551 | 56.3 | 10 | 42 | 372 | 20 | 1 |
| 5083 | 84 | 11 | 62 | 323 | 20 | 1 |
| 13380 | 84.4 | 25 | 63 | 308 | 20 | 1 |
| 17765 | 85 | 40 | 63 | 292 | 20 | 1 |
| 30027 | 135 | 37.5 | 100 | 208 | 20 | 1 |

### RNA, miRNA and Sensor-seq library preparation

Total RNA was extracted from three replicates of cultured HUVEC cells seeded on 3kpa or 30kpa PDMS using TRIzol reagent (Life Technologies) according to the manufacturer’s protocol. For mRNA libraries, total RNA was treated with DNA-free DNase (Ambion) and 500 ng of treated RNA was used to prepare Lexogen QuantSeq 3’ mRNA-Seq FWD libraries for Illumina deep-sequencing according to the manufacturer’s protocols using the i7 barcode indices. Libraries were amplified with 12 PCR cycles. miRNA libraries were prepared from 1μg of total RNA using the NEBNext^®^ Small RNA Library Kit (NEB) following the gel size selection method in manufacturer’s protocol and submitted for Illumina sequencing. For Sensor-Seq library, a customized oligonucleotide library was synthetized by Integrated DNA Technologies (IDT). The sequence of each individual oligonucleotide was obtained from the piranha analysis (see Supplementary Table 4) extending the genomic coordinate of each peak by 20 nucleotides at the 3 and 5 prime region. 97 peaks with at least 1 predicted MRE, representative of 51 CAM genes were selected. In addition, all the sensor oligonucleotides contained restriction enzyme sites, AscI and NheI, to allow for PCR base amplification and cloning. The library of oligonucleotides was resuspended in 480 μl of water, diluted 1:100 and PCR amplified as follows: in a 50 μl reaction, 10 μl 5X Phusion HotStart II HF Buffer, 1 μl 10 mM dNTP, 2.5 μl 10 mM AscI forward (5’ - GGCCATCTGGCGCGCC) and NheI reverse primers (5’-GGCCGATAAGCTAGC), 1 μl of diluted library, 1 μl DMSO, 0.5 μl Phusion HotStart II Polymerase. Cycling parameters were: 98°C for 2 min, 20 cycles of 98°C for 20 sec, 63°C for 20 sec, 72°C for 20 sec and 72°C for 2 min. PCR-amplified libraries were purified using a PCR purification kit (Qiagen) and double digested for 2 h at 37°C. Sensor-seq backbone^27^ containing a bidirectional promoter for UbC upstream of copGFP and mCherry genes, was kindly provided by Dr. Jun Lu, Yale University. After sensor backbone digestion with AscI and NheI, the MRE oligo library was cloned into the 3’UTR of mCherry. Ligation was performed in 20 μl reactions containing 50 ng vector, 3 ng insert, 2 μl 10x T4 Ligation Buffer, 2500 U T4 DNA Ligase (Promega), incubated 16 h at 16°C. After transformation into DH5α, colonies were collected, and a pool of plasmids was prepared using a Maxi prep kit (Qiagen).

Lentiviruses for expression of the CAM-MRE Sensor library were generated by transfecting Lenti-X 293T cells (10cm dishes) with packaging vectors (2.5μg VSV; 5μg pxPAX2) and the DNA vector library (7.5ug) with lipofectamine 2000 (Invitrogen). For FACS analysis additional control lentiviruses expressing GFP alone, mCherry alone, miRNA sensor lacking a MRE (Empty-Sensor plasmid, negative control), and miRNA sensor with a synthetic MRE for miR125 (miR125-Sensor plasmid, positive control) were used.

### RNA and miRNA seq data analysis

Data analysis for RNA and miRNA seq was performed using STAR alignment software and R software environment for statistical computing (http://www.r-project.org/). Total RNA and microRNA were aligned against the human genome version GRCh38 using the GENCODE 22 transcript annotation, using STAR alignment software with same parameters used for the ENDOCE project (www.encodeproject.org) After alignment, differential gene expression (DEG) of RNAs or miRNAs between ECs seeded on 30 and 3 kPa substrates was computed using the negative binomial distributions via edgeR using standard parameter Genes^62,63^. The levels (log 2 Fold Change) of CAM RNAs and SEED matching miRNAs to CAM-MRE sensors, were combined and represented in the CIRCOS plot.

### FACS and Sensor-Seq Analysis

For FACS experiments, cells were infected with low levels of the library or control lentivirus in the absence of polybrene to avoid multiply infected cells (10-20% of cells infected). After 48 h, cells were trypsinized and seeded onto soft (3kPa) or stiff (30kPa) fibronectin-coated PDMS dishes at low density (150k cells per 10cm dish) for 2 days. After 2 days on PDMS gels, cells were washed once with PBS, trypsinized, centrifuged for 5 min at 300xg, and re-suspended in PBS at 500k cells per ml immediately before FACS analysis. Infected cells were sorted on a BD FACSAria II and analyzed with FACSDiva 7. Four sorting gates were set based on the 2 control plasmids (Empty-Sensor and miR125-Sensor). The upper limit bin (Not Suppressed) was designed to contain 90% or more events/cells infected with the Empty-Sensor and less than 0.5% of events for the miR125-Sensor. Conversely, the lower two bins (Strongly Suppressed and Suppressed) were designed to contain 90% of events coming from cells infected with miR125-Sensor, in a ratio close to 3:2 (∼60% of events in Strongly Suppressed bin and ∼40% of events in Suppressed). The 3rd bin (Mildly Suppressed) was set between the Not Suppressed and the Suppressed bins. For clarity, the contour plot represents the total percentage of event in each single bins, grouped in “island” of 15% probability were shown for the Empty-Sensor, miR125-Sensor, Sensor-Library at 3 kPa and Sensor-Library at 30 kPa.

After sorting of cells into each bin, genomic DNA was isolated using the DNeasy Blood & Tissue Kit (Qiagen) and the MREs were PCR amplified using specific forward primers to barcode each bin: strongly suppressed 5’-AGCACTCGAGCTGTACAAGTAGTG-3’, suppressed 5’-TCGGAACGAGCTGTACAAGTAGTG-3’, mildly suppressed 5’-CCAGTACGAGCTGTACAAGTAGTG-3’ and not suppressed 5’-GACATCCGAGCTGTACAAGTAGTG–3’, and the reverse primer 5’-TGTAATCCAGAGGTTGATTATCG. The PCR protocol used was: in 50 μl reaction, 10 μl 5X Phusion HotStart II HF Buffer, 1 μl 10 mM dNTP, 1 μl 10 mM 5’and 3’primers, 100 ng genomic DNA, 0.5 μl Phusion HotStart II Polymerase. Cycling parameters were: 98°C for 2 min, 35 cycles of 98°C for 20 sec, 59°C for 20 sec, 72°C for 20 sec and 72°C for 5 min. Library were purified on a 1% agarose gel using gel extraction kit (Qiagen). The primers used contain barcodes for multiplexing (underline sequences) and were designed to hybridize with Illumina sequencing.

Computational analysis of Sensor-seq was performed using R software environment for statistical computing, using customized pipeline. First, the number of reads for each sensor-MRE were normalized by dividing by the total number of reads in the entire experiment. This was then multiplied by one million to get the Reads Per Million (RPM) for each sensor. To calculate the frequency of sensor-MREs in each bin, the RPM was divided by the total RPM for all 4 bins of the experiment for that sensor. This gave frequency values for each MRE in each bin at each stiffness. MREs that showed a dominant bin (with frequency values above 0.375, i.e. non-random) were compared at 3 vs. 30 kPa. In Figure 2c CAM MREs were then plotted based on the reproducible tendency to be enriched in the same bin at a given stiffness but not the other for 3 independent experiments. MREs that shifted towards a more suppressed bin on 30kPa compared to 3kPa were plotted as CAM-MRE 30kPa (bottom left of plot). MREs that that shifted towards a more suppressed bin on 3kPa compared to 30kPa were plotted as CAM-MRE3kPa (upper right of plot). RNA-seq (red) and Proteomics (green) data for each of these proteins was also plotted and linked with the miRNA-seq (for miRNAs predicted to bind to these particular MREs). The regulation of CAM-MREs by miRNAs (stars in the Figure 2c) was further validated by individual MRE mutagenesis followed by functional assays.

### Mass spectrometry sample preparation and analysis

Cells were lysed in 25 mM ammonium bicarbonate (AB) buffer containing 1.1 % sodium dodecyl sulfate (SDS, sigma), 0.3 % sodium deoxycholate (Sigma), protease inhibitor cocktail (Sigma), phosphatase inhibitor cocktails (Merck). Six 1.6 mm steel beads (Next Advance Inc.) were added to the tube and samples were homogenized with a Bullet Blender (Next Advance Inc.) at maximum speed for 2 minutes. Resulting homogenates were cleared by centrifugation (12 °C, 10000 rpm, 5 minutes). Immobilized-trypsin beads (Perfinity Biosciences) were suspended in 150 μL of digest buffer (1.33 mM CaCl_2_, Sigma, in 25 mM AB) and 50 μL of cell lysate and shaken overnight at 37 °C in a Thermocycler at 1400 rpm. The resulting digest was then reduced by the addition of 4 μL of 500 mM dithiothreitol (DTT, Sigma, in 25 mM AB; 10 min. shaking at 1400 rpm at 60 °C) and alkylated by the addition of 12 μL 500 mM iodoacetamide (Sigma, in 25 mM AB; 30 min. shaking in the dark at room temperature). Immobilized trypsin beads were removed by centrifugation at 10,000 rpm for 10 min. Supernatant containing reduced, alkylated peptides were transferred to 1.5 mL ’LoBind’ Eppendorf tubes and acidified by addition of 5 μL 10 % trifluoroacetic acid (Riedel-de Haën) in water, and cleaned by two-phase extraction (3 x addition of 200 μL ethyl acetate, Sigma, followed by vortexing and aspiration of the organic layer). Peptides were desalted, in accordance with the manufacturer’s protocol, using POROS R3 beads (Thermo Fisher) in accordance with the manufacturer’s protocol and lyophilized. Peptide concentrations (measured by Direct Detect spectrophotometer, Millipore) in injection buffer (5 % HPLC grade acetonitrile, Fisher Scientific, 0.1% trifluoroacetic acid in deionized water) were adjusted to 200 ng μL^−1^ prior to MS analysis. Digested samples were analysed by LC-MS/MS using an UltiMate^®^ 3000 Rapid Separation LC (RSLC, Dionex Corporation, Sunnyvale, CA) coupled to a Q Exactive HF (Thermo Fisher Scientific, Waltham, MA) mass spectrometer. Peptide mixtures were separated using a multistep gradient from 95% A (0.1% FA in water) and 5% B (0.1% FA in acetonitrile) to 7% B at 1 min, 18% B at 58 min, 27% B in 72 min and 60% B at 74 min at 300 nL min^−1^, using a 75 mm x 250 μm i.d. 1.7 μM CSH C18, analytical column (Waters). Peptides were selected for fragmentation automatically by data dependent analysis.

### Mass spectrometry data processing and protein quantification

Spectra from multiple samples were automatically aligned using Progenesis QI (Nonlinear Dynamics) with manual placement of vectors where necessary. Peak-picking sensitivity was set to 4/5 and all other parameters were left as defaults. Only peptides with charge between +1 to +4, with 2 or more isotopes were taken for further analysis. Filtered peptides were identified using Mascot (Matrix Science UK), by searching against the SwissProt and TREMBL mouse databases. The peptide database was modified to search for alkylated cysteine residues (monoisotopic mass change, 57.021 Da), oxidized methionine (15.995 Da), hydroxylation of asparagine, aspartic acid, proline or lysine (15.995 Da) and phosphorylation of serine, tyrosine or threonine (79.966 Da). A maximum of 2 missed cleavages was allowed. Peptide detection intensities were exported from Progenesis QI as Excel (Microsoft) spread sheets for further processing. Peptide identifications were filtered via Mascot scores so that only those with a Benjamini-Hochberg FDR < 0.05 remained. Raw ion intensities from peptides belonging to proteins with fewer than 2 unique (by sequence) peptides per protein in the dataset were excluded from quantification. Remaining intensities were logged and normalized by the median logged peptide intensity. Missing values were assumed as missing due to low abundance, an assumption others have shown is justified^64^. Imputation was performed at the peptide level following normalization using a method similar to that employed by Perseus^65^ whereby missing values were imputed randomly from a normal distribution centered on the apparent limit of detection for this experiment. The limit of detection in this instance was determined by taking the mean of all minimum logged peptide intensities and down-shifting it by 1.6σ, where σ is the standard deviation of minimum logged peptide intensities. The width of this normal distribution was set to 0.3σ as described in^65^. Fold-change differences in the quantity of proteins detected in different time-points were calculated by fitting a mixed-effects linear regression model for each protein with Huber weighting of residuals as described in^64^using the fitglme Matlab (The MathWorks, USA) function with the formula:

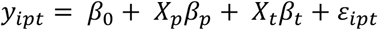

Where *y_iptb_* represents the log_2_(intensity) of peptide *p* belonging to protein i, under experimental treatment *t. βs* represent effect sizes for the indicated coefficients. *Peptide* effects were considered as random effects whereas *treatment* was considered as a fixed effect. ß_0_ denotes the intercept term and ε_*PT*_ denotes residual variance. Standard error estimates were adjusted with Empirical Bayes variance correction according to ^66^. Conditions were compared with Student’s t-tests with Benjamini-Hochberg correction for false positives.

### Cells Immunostaining and Quantification

Cells seeded on fibronectin-coated PDMS were fixed with 4% paraformaldehyde (Electron Microscopy Sciences) in PBS. Cells were then washed and permeabilized with 0.05% Triton X-100 in PBS supplemented with 320 mM sucrose and 6 mM MgCl_2_. Cells were washed 3 times with PBS and blocked for 30 minutes with 1% BSA in PBS. Cells were incubated overnight at 4°C with anti-YAP antibody (1:200, no sc-101199, Santa Cruz Biotechnology), anti-RhoB (1:250, 19HCLC, Thermo Fisher), anti-Vinculin (1:200, V9131, Sigma-Aldrich), anti-STMN1 (1:200, ab52630, Abcam), anti-CTGF (1:200, ab6992, Abcam) and anti-paxillin (1:800, RabMAb Y113, ab32084, Abcam) diluted in 1% BSA in PBS. Cells were washed 3 times with PBS and incubated at room temperature for 1 hour with secondary antibodies (alexa-488 anti-rabbit, alexa-647 anti-mouse, 1:1000, Molecular Probes) and alexa-565 conjugated phalloidin (1:1000, molecular probes). Cells were washed again 3 times with PBS and mounted with DAPI in Fluoromout-G (SouthernBiotech). Cell areas were quantified using ImageJ by background subtracting, thresholding to generate cell masks, and using the analyze particles function (n=51-150 cells / group, experiment replicated 3 times for each cell type). YAP staining was quantified by taking the average nuclear YAP signal (in the area of the DAPI stain), divided by the average cytoplasmic YAP signal (in the area of the non-nuclear cell mask). Focal adhesions were analyzed using the focal adhesion analysis server ^67^ with the minimum adhesion size set to 0.5 μm^2^ and the default settings for only static properties.

### Traction Force Microscopy

PDMS TFM substrates were fabricated as described ^68^. Briefly, cover-glass bottom dishes were spin-coated to obtain a ∼40μm thick layer of polydimethylsiloxane (PDMS; Sylgard 184 by Down Corning mixed at various B/C ratios, 67:1 3kPa, 40:1 30kPa) and cured at 70°C for 3 hours. Gels were then treated with 3-aminopropyl trimethoxysilane for 5 min and incubated for 10 min at room temperature with 40nm Alexa Fluor 647 beads (Molecular Probes) suspended in a 100μg/ml solution of 1-Ethyl-3-(3-dimethylaminopropyl) carbodiimide (Sigma) in water to covalently link the beads to the gel surface. Elastic moduli for each batch was measured using a microfluidic device as described ^68^ and is reported as the Young’s modulus (E).

TFM gels were coated with fibronectin (10μg/ml) in PBS overnight at 4°C and washed 3 times with PBS. HUVECs and HDFs were seeded on the gels in EGM or low serum fibroblast growth medium, respectively, 24h before analysis at low density (∼3000 cells per cm^2^). Cells and florescent beads were imaged on a spinning disk confocal microscope (UltraVIEW VoX, Perkins Elmer) attached to a Nikon A-1 microscope equipped with a temperature and CO_2_ controlled incubation chamber and 60x 1.4NA lens. Florescent images of Alexa Fluor 647 beads and DIC images of cells were acquired before and after cell lysis with 0.05% SDS. Images were drift corrected and bead displacements were quantified using a previously developed open source traction force microscopy software in MATLAB 2015a ^69^. Force fields and traction stresses were calculated using FTTC force reconstruction with the regularization parameter set to 0.007. Total force per cell was calculated as the average stress under the cell multiplied by the area.

### 3D matrix constructs

A method to generate three-dimensional cell-derived uniaxial matrix constructs (3D matrix constructs) based on the “tendon construct” developed by Karl Kadler’s group was used^70^. Six well plates were coated with a 2 mm layer of SYLGARD 184 and incubated overnight at 65°C to induce polymerization. After cooling, the (hydrophobic) SYLGARD layer was incubated 15 minutes with Pluronic^®^ F-127. Custom-made rectangular Teflon molds (15×10×2mm) were sterilized with Virkon (10 minutes) then Ethanol 70% (15 minutes). Inside the molds, two 8mm segments of size 0 silk sutures were pinned to the PDMS using insect pins exactly 10mm apart. Inside each mold, we added 12 μL of thrombin stock solution (200U/mL, Sigma). Primary fibroblasts were detached with 0.05% trypsin, centrifuged at 1800 rpm and counted. For each matrix construct, 2×105 cells were resuspended in 300 μL of DMEM containing 8mg/mL fibrinogen (Sigma) and 0.2 mM of L-ascorbate-2-phosphate. Cell suspensions were injected inside the molds and placed at 37°C for 15 min in incubator for polymerization. After polymerization, the Teflon mold was removed and one additional insect pin added to maintain the suture thread. Matrix constructs were cultured with DMEM/F12 supplemented with 10% fetal bovine serum (Sigma), Penicillin (100U/mL), Streptomycin (100 ug/mL, Gibco), 1% L-glutamine and 0.2mM of L-ascorbate-2-phosphate. The 3D matrix constructs were cultured for 5 days and the culture medium changed every other day. After 5 days, photographs of the constructs were taken with a Nikon reflex camera equipped with a 50mm macro-objective at a focal distance of 1:1. Constructs diameter was obtained by averaging the diameter at 3 different locations (each extremity and the middle).

### Immunostaining matrix constructs

Matrix constructs were rinsed in cold PBS and fixed overnight at 4°C in 4% formaldehyde (Pierce 16% formaldehyde, methanol free) in PBS. Fixation constructs were dehydrated, embedded in paraffin and 5 μm transverse sections cut with a Leica microtome. For immunostaining, we performed a rehydration protocol followed by antigen retrieval for 30 minutes at 96°C in a citrate buffer (pH 6). Sections were blocked with Odyssey PBS blocking buffer (LiCor biosciences) for 1 hour and incubated overnight with primary antibodies diluted in blocking buffer: vimentin (1/400, Cell Signaling), phospho-myosin light chain (1/400, Abcam). After extensive rinsing in PBS Tween 0.1%, slides were incubated with AlexaFluor 647 anti-rabbit secondary antibody (1/500, ThermoScientific) for 1 hour at room temperature, thoroughly washed with PBS Tween 0.1%, and slides mounted in FluoromountG-DAPI (Southern Biotech). Slides were imaged with an Olympus slide scanner microscope equipped with a 20x objective.

### Zebrafish Fin Fold Regeneration

Zebrafish were raised and maintained at 28.5°C using standard methods and according to protocols approved by Yale University Institutional Animal Care and Use Committee (# 2015-11473). Wild-type (AB) and mz *ago2* -/- mutants^38^ were used. To generate the ctgfa MRE mutant, zebrafish AB were injected with 125 ng/μl *Cas9* mRNA and 75 ng/μl gRNAs, designed as previously described^59^. The gRNA sequence used to target the conserved MRE within the 3’UTR human *CTGF* gene was (5’-GGTGAAAACATGTAACATTT-3’). Genomic DNA was isolated from a clutch of 15 injected and uninjected control embryos at 24 hpf using the Qiagen DNeasy Blood and Tissue Kit. Genomic DNA (250 ng) and the Phusion HotStart II Kit (ThermoFisher) used to PCR amplify an approximately 300 bp region surrounding the intended MRE target (Fwd: 5’-TTGGGAAAGAGCCAGTATCC-3’, Rev: 5’-TGGTGCCATTATTGTGTGGT-3’). T7 Endonuclease I assay was used to detect mutations as described in the manufacturer’s protocol (New England BioLabs). PCR and T7 products were run on 3% agarose gels to verify the occurrence of indels in the MRE sequence. The remaining embryos were grown to 48 hpf and used for the fin fold regeneration experiments (see below).

The zebrafish miR-124 and ctgfa sensor assay and mRNA injection were performed as described^59^. For the fin fold regeneration assay, we use 14 AB fish, 14 mz ago2 mutant -/- embryos and 15 mz ago2 -/- fish injected at the one cell stage with 200 pg of in vitro transcribed mRNA encoding the human AGO2 protein. At 2 days post fertilization (dpf), the fin fold was cut at the edge of the fin using a 25G needle. Bright filed images were captured at 0.5, 2, 4, 24, 48 and 72 hours post amputation (hpa) using a Leica M165 FC stereomicroscope and Leica Application Suite V4 software. The length of the fins over time was measured using FIJI-ImageJ ^70,71^ and normalized for the length of the fin before cutting. Analysis and graph were generated using Graphpad Prism7 statistical software.

### Zebrafish immunofluorescence assay

For the fluorescent images: 20 embryos for each genotype (AB, Ago2 mutant (-/-) and ctgfa 3’ UTR mutant) were cut and then, at 0.5, 2, 4, 24 hpa were fixed in PFA 4% overnight at 4°C. Embryos were washed 4-5 times with PBS 0.1%Tween, then incubated 2 hours in blocking solution (0.8% Triton-X, 10% normal goat serum, 1% BSA, 0.01% sodium azide in PBS Tween). Zebrafish were stained following the protocol as in^13^ using the primary antibody mouse anti-Phospho-Myosin Light Chain 2 (1:200; Cell Signaling), mouse anti-Proliferating Cell Nuclear Antigen (1:200; PCNA, Dako), rabbit anti-Fibronectin (1:200; Cell Signaling), DAPI (1:1000; Sigma), rabbit anti-Connective Tissue Growth Factor A (1:150; Abcam), and mouse anti-YAP (1:200; Santa Cruz Biotechnology) and the secondary antibody Alexa Fluor 488 anti-mouse (1:250, ThermoFisher) and Alexa Fluor 596 anti-rabbit (1:250, ThermoFisher). After staining, images were captured using a Leica Microsystems SP5 confocal microscope using a 40X objective. Max projections were generated with the Leica application suite or Perkin Elmer Volocity software. Intensity was quantified using FIJI-Image. For each protein staining, a line profile of 80 µm in diameter within the wound was calculated for the first 50 μm from the fin fold edge. The intensity profile of 4 to 6 fish was calculated using R statistical software. To determine the ratio of Nuclear and Cytosolic YAP in before and during the fin fold regeneration confocal images were split by channel and a threshold was used on the DAPI channel to generate a binary mask for the nuclei. Using the Image Calculator function of ImageJ, the binary mask was subtracted from the YAP channel to generate the cytosolic YAP image. After inverting the binary mask and subtracting it from the YAP channel, the nuclear YAP image was generated. Each were measured and normalized to the area. For TUNEL assay to detect apoptotic cells embryos were fixed in 4% PFA overnight and stored in 100% methanol at −20 °C. The TUNEL assay was performed using the ApopTag Red *In situ* Apoptosis Detection kit (Millipore).

### Atomic Force Microscopy

Live zebrafish embryos (48 hpf) were anesthetized using 1x tricaine in egg water and mounted on PDMS gels. The tips of the fish tails were probed using a DNP-10 D tip (Bruker, nominal stiffness ∼0.06 N/m) on a Bruker Dimension FastScan AFM immersed in egg water containing 1x tricaine. Probe deflection sensitivity was calibrated by taking indentation curves on glass and the nominal tip stiffness was calibrated by thermal tuning (assuming a simple harmonic oscillator in water). Force vs. deflection curves were collected for a ramp size of 1.5μm at a rate of 750 nm/s for at least 2 locations per fish, with 10-11 fish measured per group (n=27-49 total measurements per group). The first 600nm of the extension curves were fit with NanoScope Analysis Software version 1.5 (Bruker) assuming a Poisson’s ratio of 0.5 and using the Sneddon fit model ^73^.

### Statistical Analyses

All the of statistical analysis were performed using Prism version 7.01 (GraphPad) and R software environment for statistical computing, except for the peak identification, which used piranha software^52^ to measure the significance of read coverage height for each mapped position using the zero-truncated negative binomial model (ZTNB). To confirm changes in cell area, focal adhesion number, YAP localization and traction force generation, t-tests were performed using Prism to assess the change in the mean between Wild-type and AGO2 or MRE CRISPR/Cas9 mutant cells. These data sets contained more than 30 individual measurements for each condition and showed a log-normal distribution. For the *in vivo* analysis of zebrafish wild-type and Ago2 mutants, changes in fin fold tissue were analyzed using t-tests; fin fold regeneration was analyzed via 2 way ANOVA and Sidak’s multiple comparisons test using Prism. The distribution of fluorophore intensity within 20 μm from the edge of the wound was calculated using Prism 4th order smoothing with 2 neighbors.

### Data Availability

The accession number for all the sequence reads reported in this paper are: HITS-CLIP: GEO: GSE99686; RNA-seq and sRNA-seq for HUVEC cells at 3 kPa and 30 kPa: GEO: GSE110211; Proteomics results are reported as excel file Supplementary_table2

**Supplementary Figure 1.**
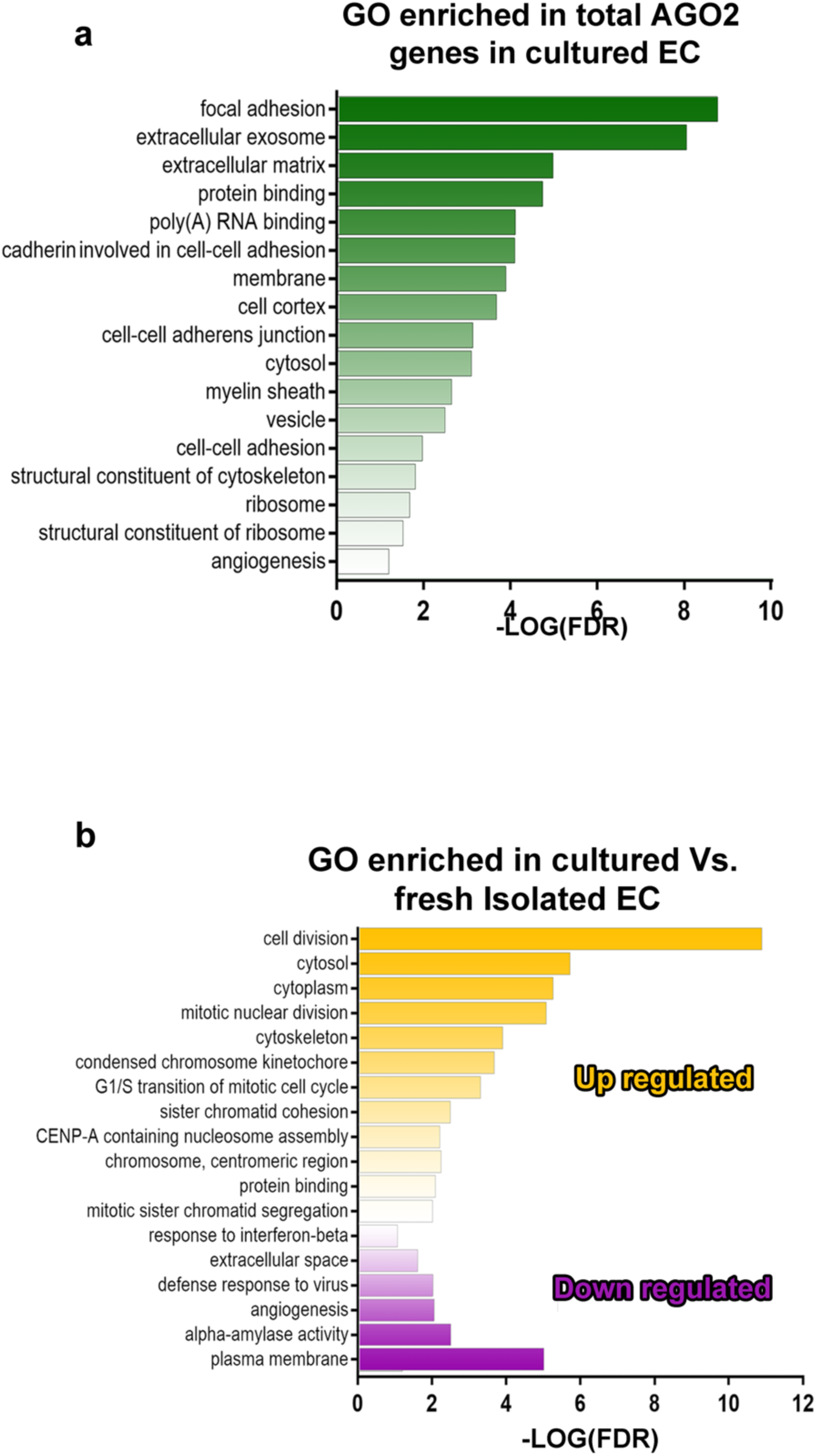
AGO2-peaks are positioned preferentially on cytoskeleton-adhesion-matrix (CAM) genes. (a)Bar graph of the Log(FDR) for the significant enriched Gene Ontology terms resulting from the HITS-CLIP assay and identified by DAVID software. **(b)** Bar graph of the Log(FDR) for the significant enriched Gene Ontology terms resulting from the microarray analysis of endothelial cells in culture versus freshly isolated.

**Supplementary Figure 2.**
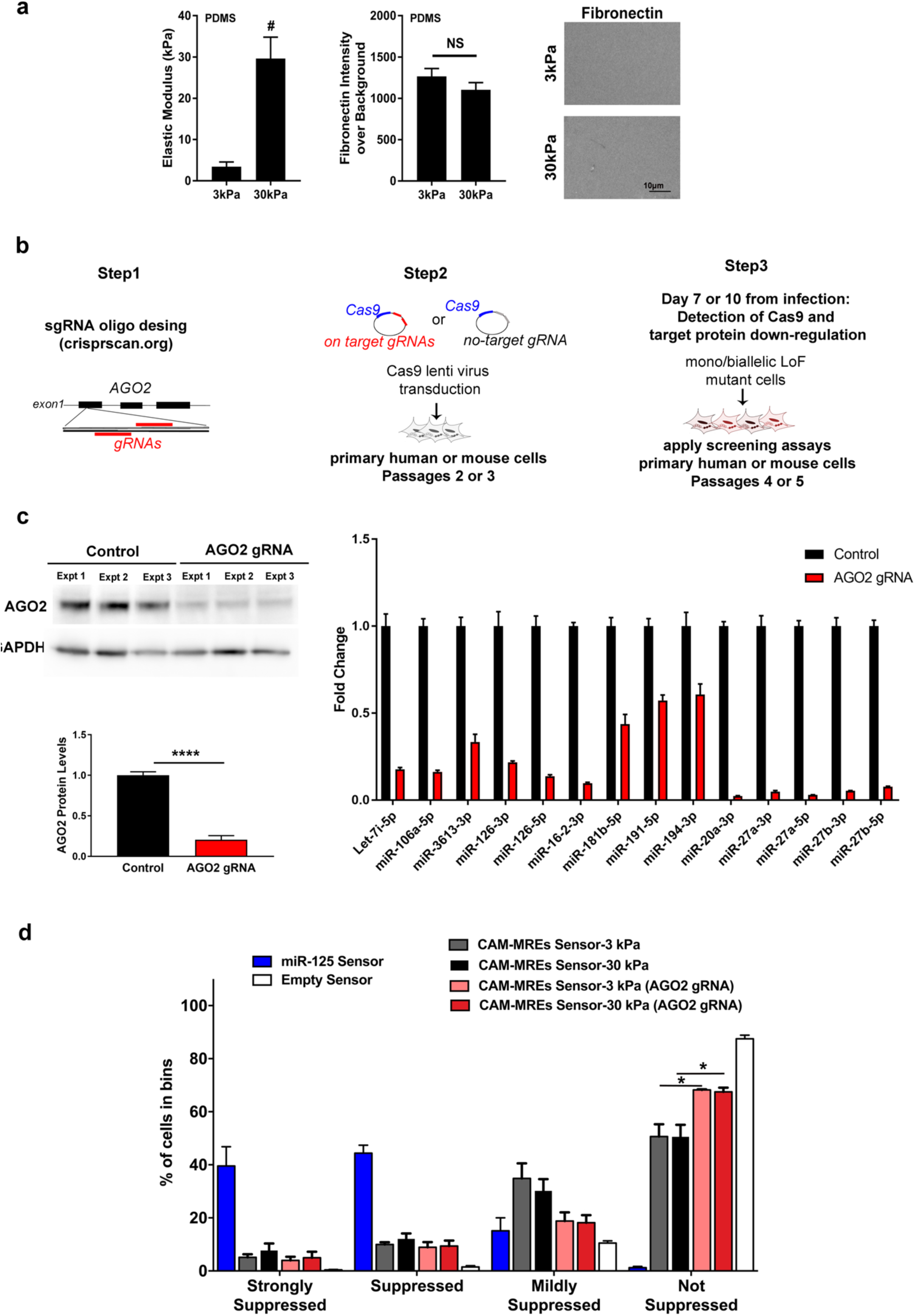
Validation of CRISPR/Cas9 targeting of AGO2 in HUVECs. (a)Quantification of elastic modulus of PDMS gels by compression testing on an instron 5848 (a, mean ± SEM, n=2 gels per condition). Gels were compressed with a cylindrical indenter to 10% strain at 0.1%/s and allowed to stress-relax for 150 seconds, modulus was measured at equilibrium. Quantification of Fibronectin staining intensity on the two PDMS gels with representative images showing uniform Fibronectin deposition on the gel surface (mean ± SEM, n=14 fields of view per stiffness). **(b)** Schematics showing the experimental approach to generate AGO2 CRISPR/Cas9 mutant cells in primary human and mouse cells. **(c)** Representative Western blot of three independent replicates showing reduced AGO2 levels in HUVECs with a lentivirus co-expressing Cas9 and *AGO2* gRNAs, or a no-target gRNA (control, has no homology to any known mammalian gene). The bar graphs show quantification of the proteins normalized to GAPDH (mean ± SEM; n = 3). qRT-PCR analysis of endothelial specific miRNAs in HUVECs infected with Cas9 and gRNA targeting *AGO2* or *DROSHA* and no-target gRNA (control) at 7 days post infection (dpi). Results are shown normalized to controls (mean ± SEM; n = 3).**(c)** Bar plot representing distribution obtained upon FACS analysis of 10,000 wild-type or AGO2 gRNA mutant HUVECs infected with CAM MRE Sensors and treated as indicated (mean ± SEM; n = 3, * p<0.05)., mCherry/GFP ratio of CAM MRE Sensors increase in AGO2 mutant ECs, as the cell distribution is shifted significantly toward the Empty-Sensor control bin in both 3 and 30 kPa stiffness conditions.

**Supplementary Figure 3.**
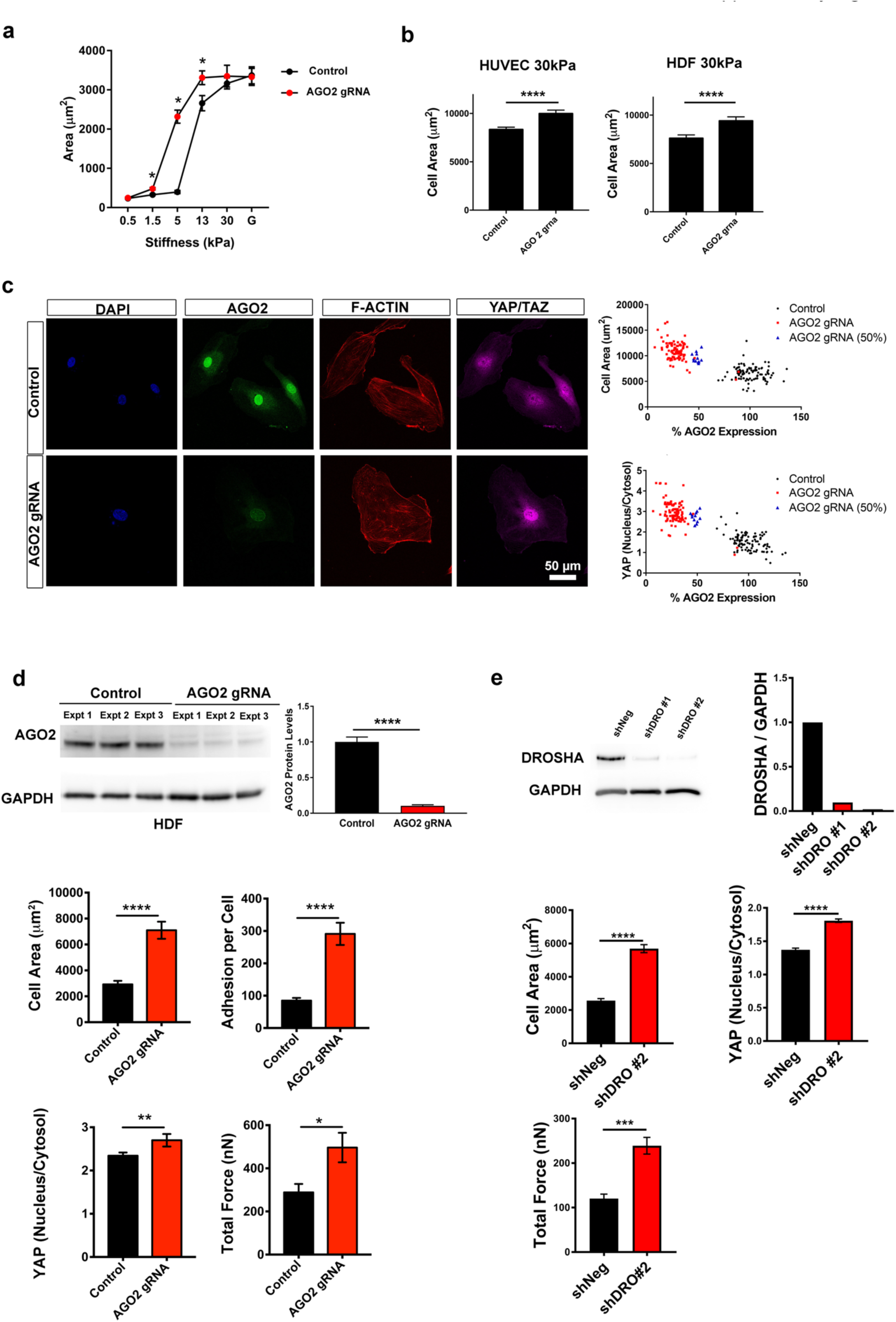
Phenotype in cells with CRISPR/Cas9 targeting of AGO2 and DROSHA. (a)Cell spread area and nuclear/cytoplasmic YAP ratio using fibronectin coated polyacrylamide gels over a wider range of stiffness display results consistent with results from PDMS gel experiments (Fig. 3a) (mean ± SEM, n=32-79 cells / group). **(b)** Bar plot representing HUVEC and HDF cell area for cells infected with AGO2 gRNA and non-targeting control seeded on fibronectin coated 30kPa PDMS gels (mean ± SEM, **** p<0.0001, HUVEC n=239-393, HDF n=201-273). **(c)** Left: Representative immunofluorescence images of HUVECs after infection with pLentiCRISPR virus directed at *AGO2* or a non-targeting control seeded on fibronectin coated 3kPa PDMS gels. Right: Quantification of AGO2 expression in ∼100 cells. Intensity of the staining in each cell is normalized for the average intensity of all AGO2 cells and transformed in %. Cell area based on F-ACTIN staining or nuclear to cytoplasmic ratio of YAP/TAZ are plotted as individual value per cell. Highlighted in red cells infected with gRNA targeting *AGO2,* in black with no-target gRNA and in blue with gRNA targeting *AGO2* expressing ∼50% expression of AGO2 compared to control cells. **(d)** Top, representative Western blot of three independent replicates showing reduced levels of AGO2 in human dermal fibroblasts (HDFs) infected with Cas9 and *AGO2* gRNAs or a no-target gRNA (control) at 7 days post infection (dpi). Bar plot indicates Western blot quantification of the AGO2 protein normalized for GAPDH at 7 dpi (mean ± SEM; n = 3). Bottom, quantifications of HDFs after infection with pLentiCRISPR virus directed at AGO2 or a non-targeting control seeded on fibronectin coated 3kPa PDMS gels. Bar plot shows HDF cell area (n=229-309 cells/group) based on phalloidin staining, number of paxillin adhesions per cell (n=34-58 cells/group), and nuclear to cytoplasmic ratio of YAP/TAZ (n=34-58 cells/group). (mean ± SEM, *p<0.05, **p<0.01,****p<0.0001). Single cell maps of average traction stress and quantification of total force per cell (mean +/-SEM, n=20-21 cells per group, * p<0.05). **(e)** Western blot for DROSHA after knockdown with shRNA delivered via lentivirus (5 days post infection, pSMART Dharmacon) with quantification on 3kPa PDMS gels of cell spread area, YAP nuclear localization (mean +/-SEM, n=138-156 cells / group, **** p<0.0001), and total force per cell (mean +/-SEM, n=21 cells / group, *** p<0.001).

**Supplementary Figure 4.**
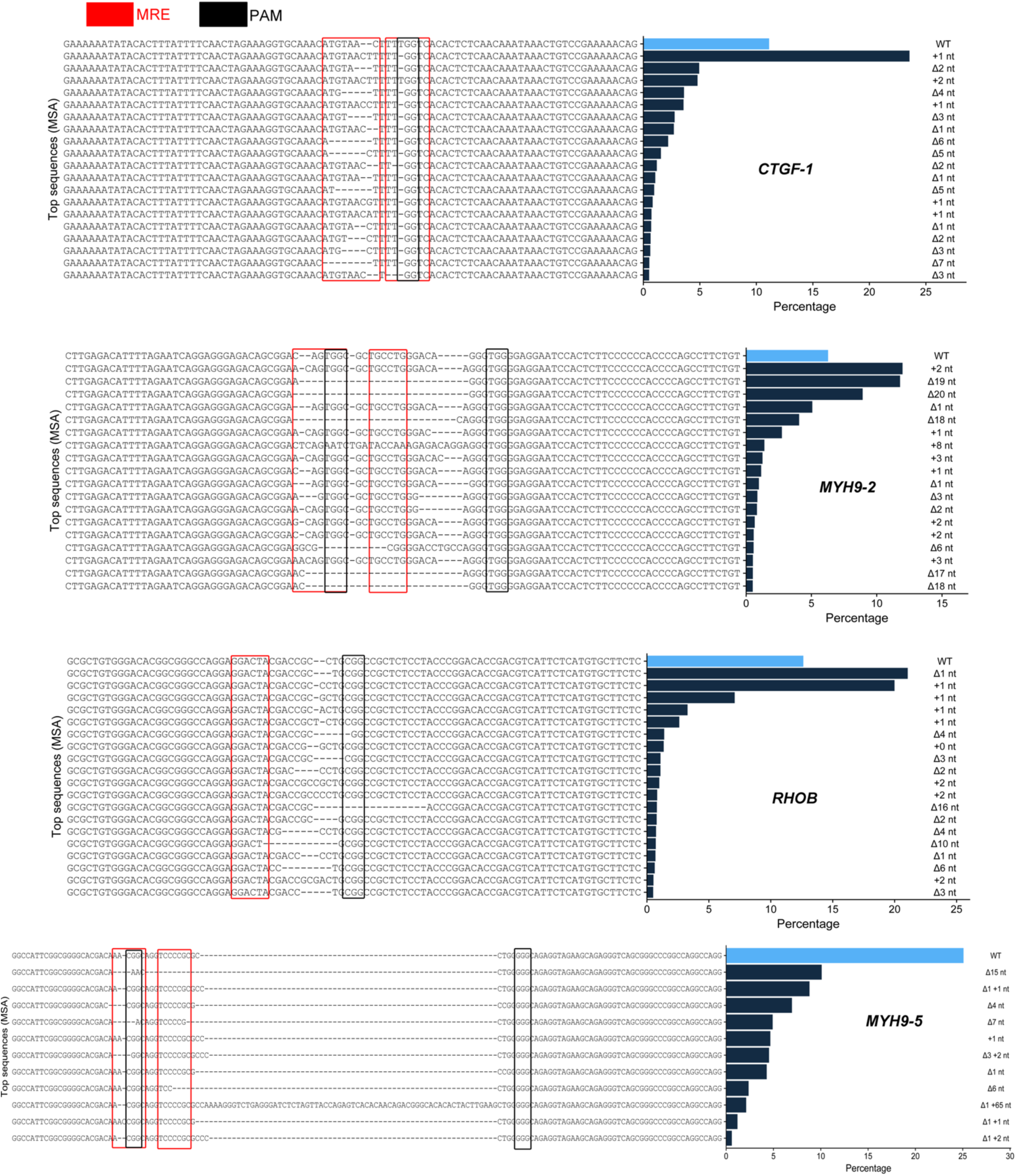
Sequences of CAM MREs resulting from CRISPR/Cas9 mutagenesis. MiSeq 2×250 analysis of PCR amplicons derived from genomic DNA of HUVEC mutated with the indicated CAM MRE gRNAs. MREs are highlighted with red boxes, PAMs are highlighted with black boxes. Bar blot represent the % wild-type and mutated CAM 3UTR sequences. Only mutations with a frequency > of 2% are represented. MSA= Multiple Sequence Alignment; nt= nucleotide. Numbers indicate the mutations as nt inserted (+) or deleted (Δ).

**Supplementary Figure 5.**
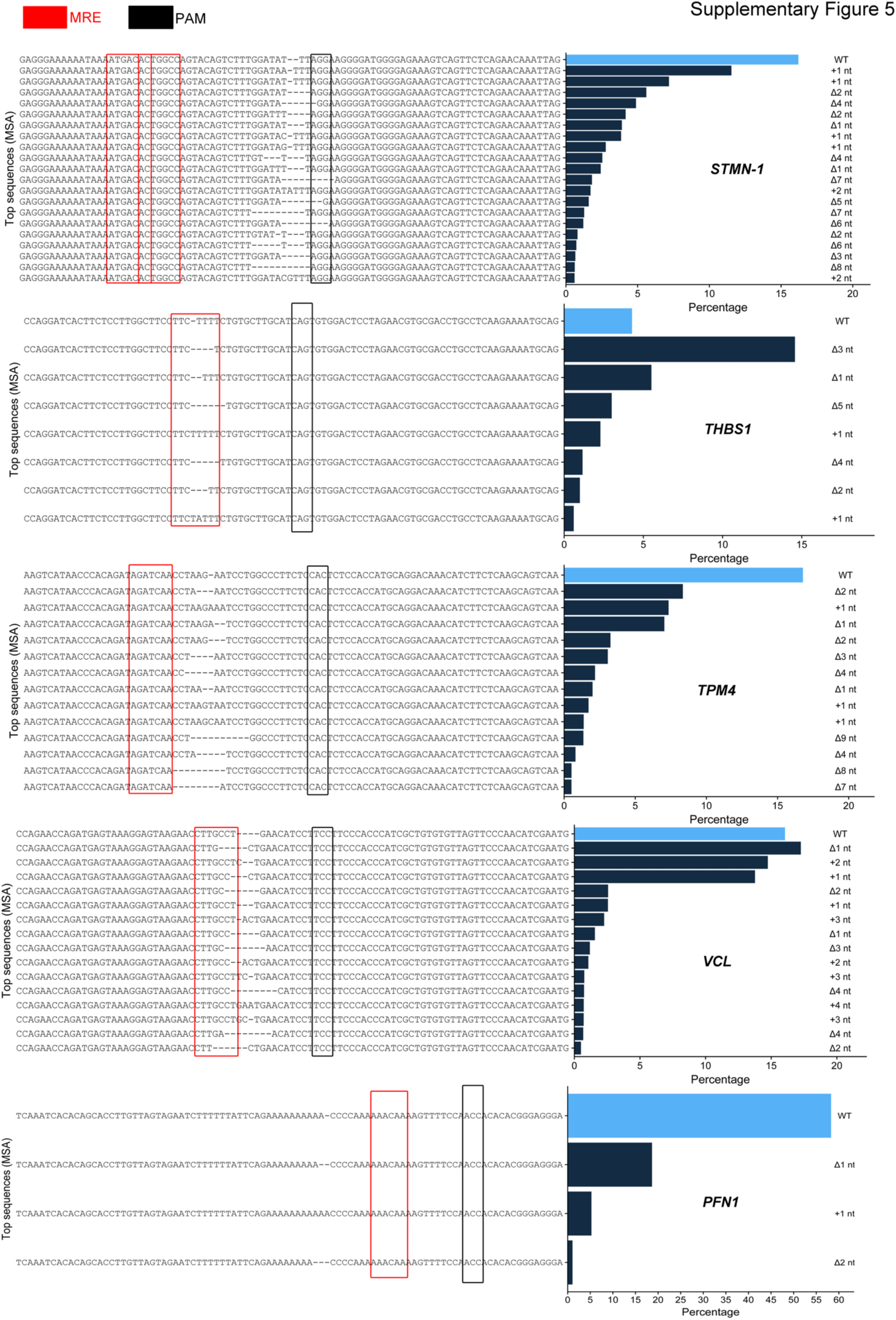
Sequences of CAM MREs resulting from CRISPR/Cas9 mutagenesis. MiSeq 2×250 analysis of PCR amplicons derived from genomic DNA of HUVEC mutated with the indicated CAM MRE gRNAs. MREs are highlighted with red boxes, PAMs are highlighted with black boxes. Bar blot represent the % wild-type and mutated CAM 3UTR sequences. Only mutations with a frequency > of 2% are represented. MSA= Multiple Sequence Alignment; nt= nucleotide. Numbers indicate the mutations as nt inserted (+) or deleted (Δ).

**Supplementary Figure 6.**
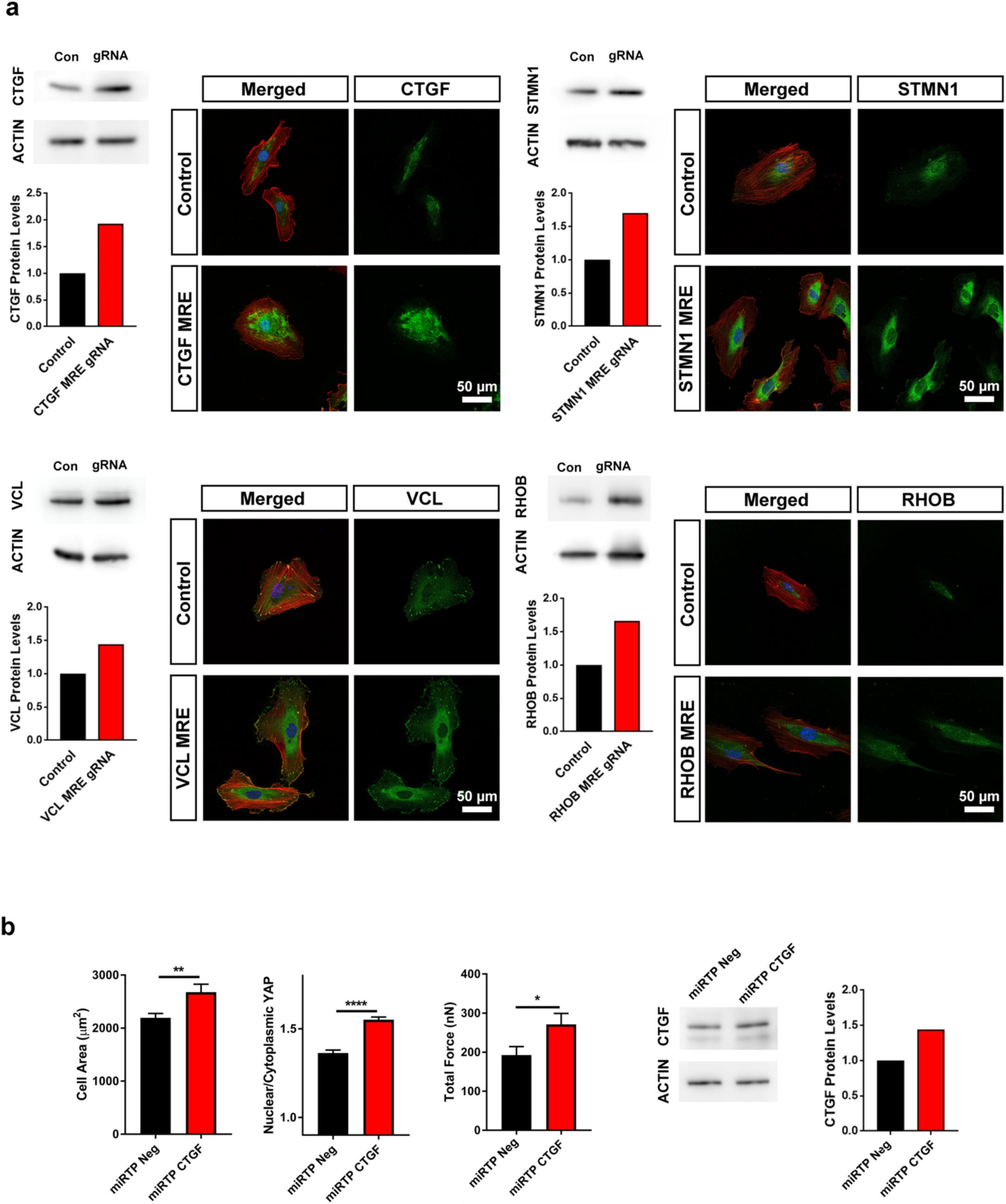
Effects of CAM MRE mutations in HUVECs. (a)Representative Western blot and immunofluorescence images showing the upregulation of the target CAM protein in human endothelial cells (HUVECs) infected with Cas9 and gRNA targeting specific CAM MREs and no-target gRNA (control) at 7 days post infection (dpi). Bar plot indicates Western blot quantification of the CAM protein normalized to β-ACTIN at 7 dpi. **(b)** Bar plots representing the quantification of mechanical parameters as in Fig. 3a of HUVECs transfected with miRTP (miRNA target protector) directed at CTGF 3UTR for 4 days. miRTP is a single-stranded, modified RNA oligonucleotide that blocks a miRNA interaction with an individual MRE. Negative control target protector (QIAGEN) has no homology to any known mammalian gene (mean ± SEM, Area and YAP: n=99-101 cells / group, TFM: n=13-15 cells/ group). *p<0.05, **p<0.01,****p<0.0001.

**Supplementary Figure 7.**
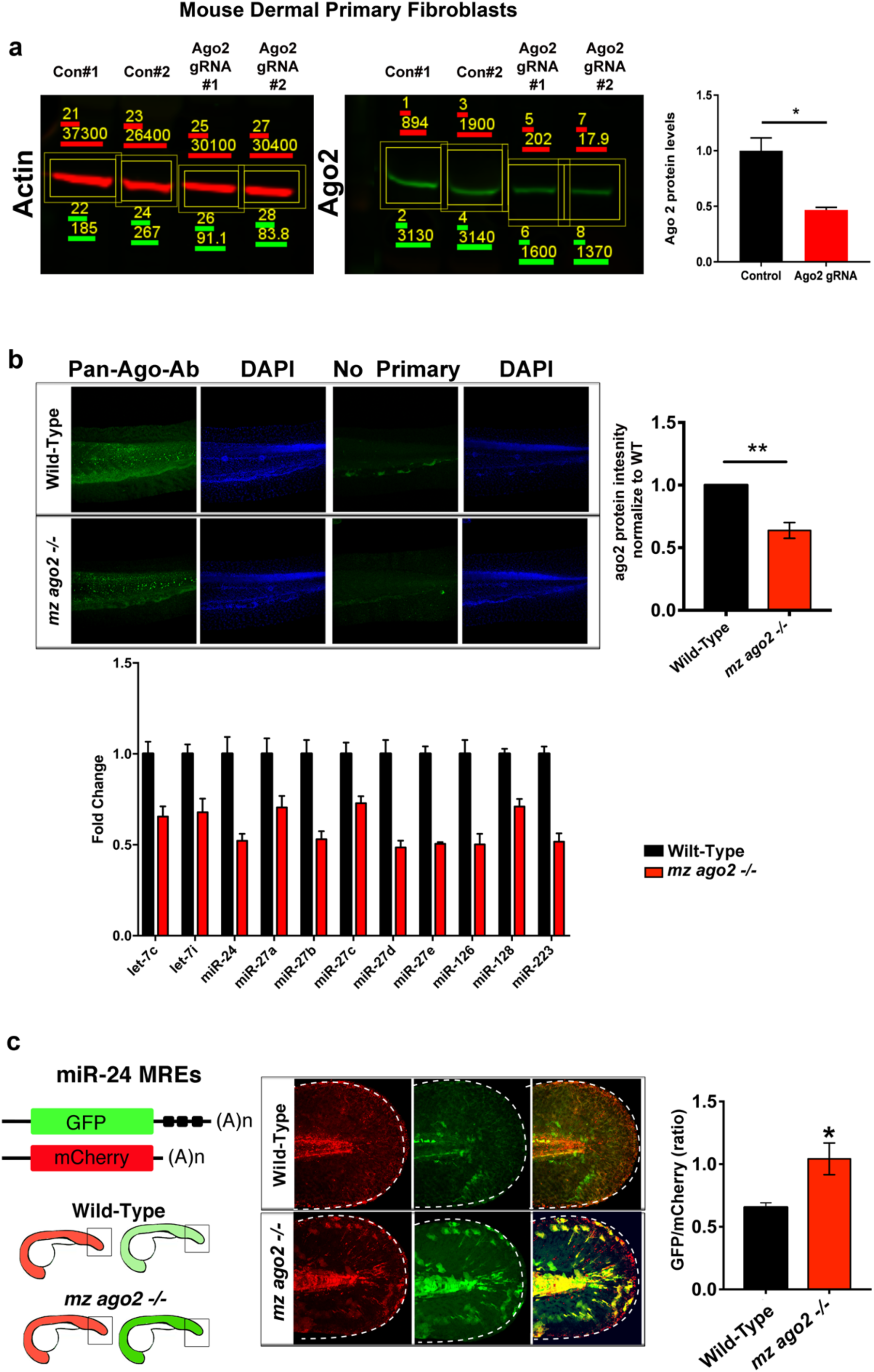
Reduced levels of Ago2 in Mouse Dermal Primary fibroblasts, and in *mz ago2 -/-* zebrafish embryos. (a) Representative blot showing reduced Ago2 levels in mouse skin fibroblast for 3D culture assay infected with gRNAs targeting Ago2 and no-target gRNA (control, has no homology to any known mammalian gene). Blots were obtained with Odyssey system from Licor to quantify near-IR fluorescence emitted by the secondary antibodies. Bar plots indicate densitometry measurements the AGO2 protein normalized for β-Actin at 7 dpi (mean ± SEM, n=2). **(b)** Top, Confocal lateral view of whole mount zebrafish embryos at 48 hours post fertilization (hpf). Wild-type and ago2 maternal zygotic homozygous mutant *(mz ago2-/-)* were stained with the Ab-panAGO2-2A8 and secondary only as a control of the total staining background. Bar plots indicate average of Ago2 fluorescence intensity normalized for the DAPI staining for each genotype (mean ± SEM, n=10 embryos / group). Bottom, qPCR representing miRNAs level in wild-type vs *mz ago2 -/-* at 72 hpf (mean ± SEM, n=3). **(c)** Right panel, schematics representing the experimental procedure to test defective miRNA-mRNA interaction in the *ago2-/-* fin fold model. *in vitro* transcribed RNAs encoding for mCherry control and GFP coding sequence upstream of three perfect MREs for miR-24, a miRNA expressed in epidermis^30^. 25 picograms (pg) of each mRNA was co-injected into wild-type and ago2-/- zebrafish embryos at the one cell stage post fertilization. Left panel, confocal whole mount image of 48 hpf embryos expressing GFP and mCherry (head is to the left). GFP and mCherry pixel intensities were quantified and the GFP/mCherry ratio was plotted for each genotype (mean ± SEM, n∼5 embryo / group). *p<0.05, **p<0.001

**Supplementary Figure 8.**
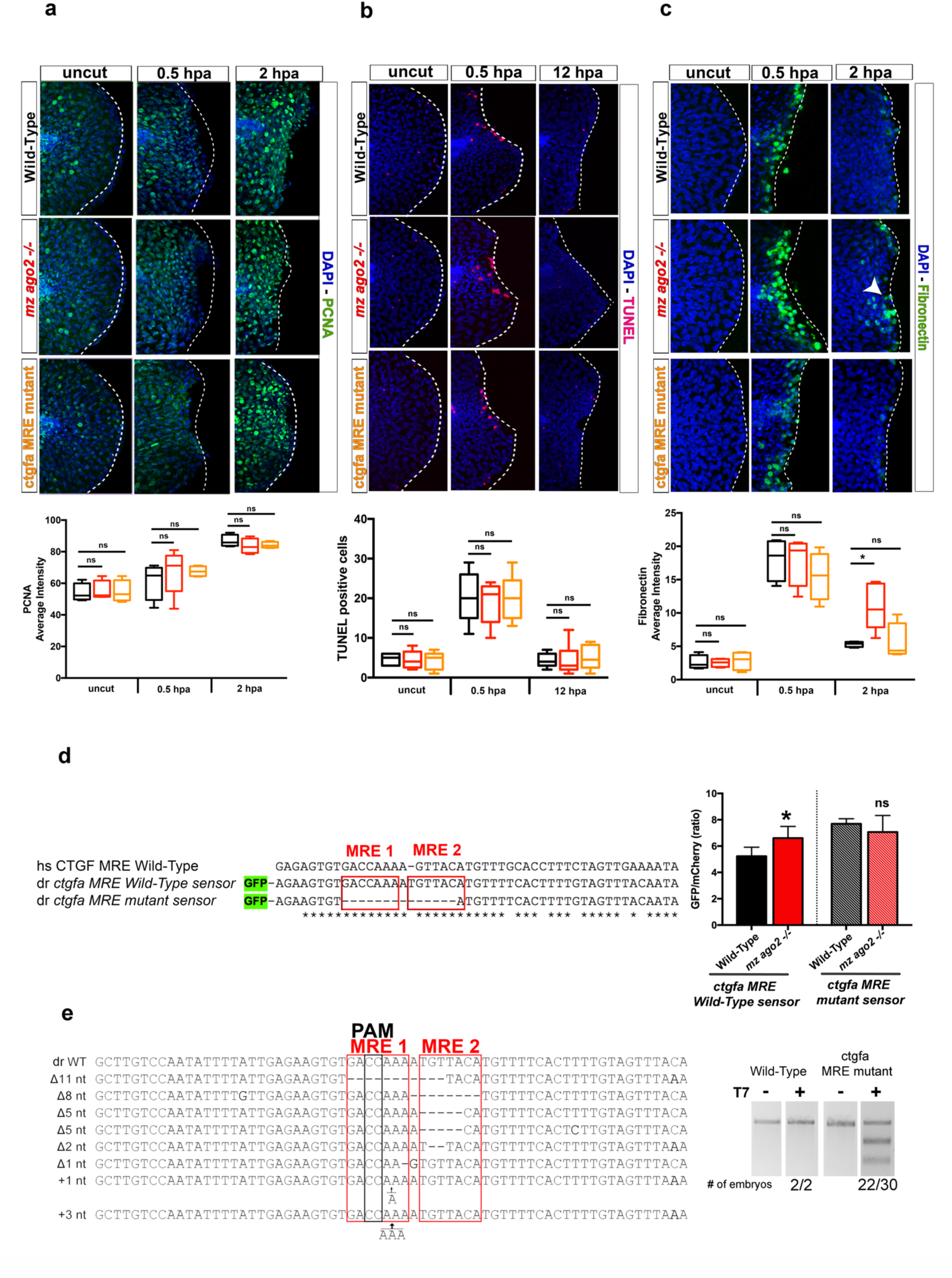
Fin fold regeneration model in zebrafish wild-type and mutant embryos. (a-c) Top, Confocal lateral view of whole mount zebrafish embryos treated as indicated. White dotted line shows the edge of the wound. Bottom, quantification of each whole mount stained embryos as indicated. Pixel matrix of intensity was reconstructed for 50 μm within the wound edge to the fish body. Pixel mean ± SEM was calculated for 4-6 fish within each group. *p<0.05, n.s.= non-significant **(d)** Alignment of human and zebrafish *ctgfa 3UTR* sequence containing the conserved MREs. This 3’ UTR sequence was used to generate a wilt-type and mutated miRNA sensor vector (as above). An in vitro transcribed mRNAs encoding for a GFP reporter was under the post-transcriptional regulation of ctgfa 3UTR (wild-type), a mutated version lacking the MREa sequence and co-injected with an mCherry mRNA with no 3UTR regulation as negative control. 75 picograms (pg) of each mRNA was co-injected into wild-type and mz ago2-/- zebrafish embryos at the one cell stage post fertilization. GFP and mCherry pixel intensities were quantified and the GFP/mCherry ratio was plotted for each genotype (mean ± SEM, n∼10 embryo / group *p<0.05). **(e)** Sequence alignment of the ctgfa 3UTR of zebrafish wilt-type and ctgfa MRE embryos injected with Cas9 and the gRNAs which PAM region was highlighted in black. Mutations are represented as nucleotide (nt) inserted (+) or deleted (Δ) and were cloned from individual embryos 24 hours post injection. Red boxes represent the MREs targeted. Representative agarose gel for T7 Endonuclease I assay for Wild-Type and ctgfa MRE mutant zebrafish. hpa= hour post amputation. ns= non significant.

**Supplementary Figure 9.**
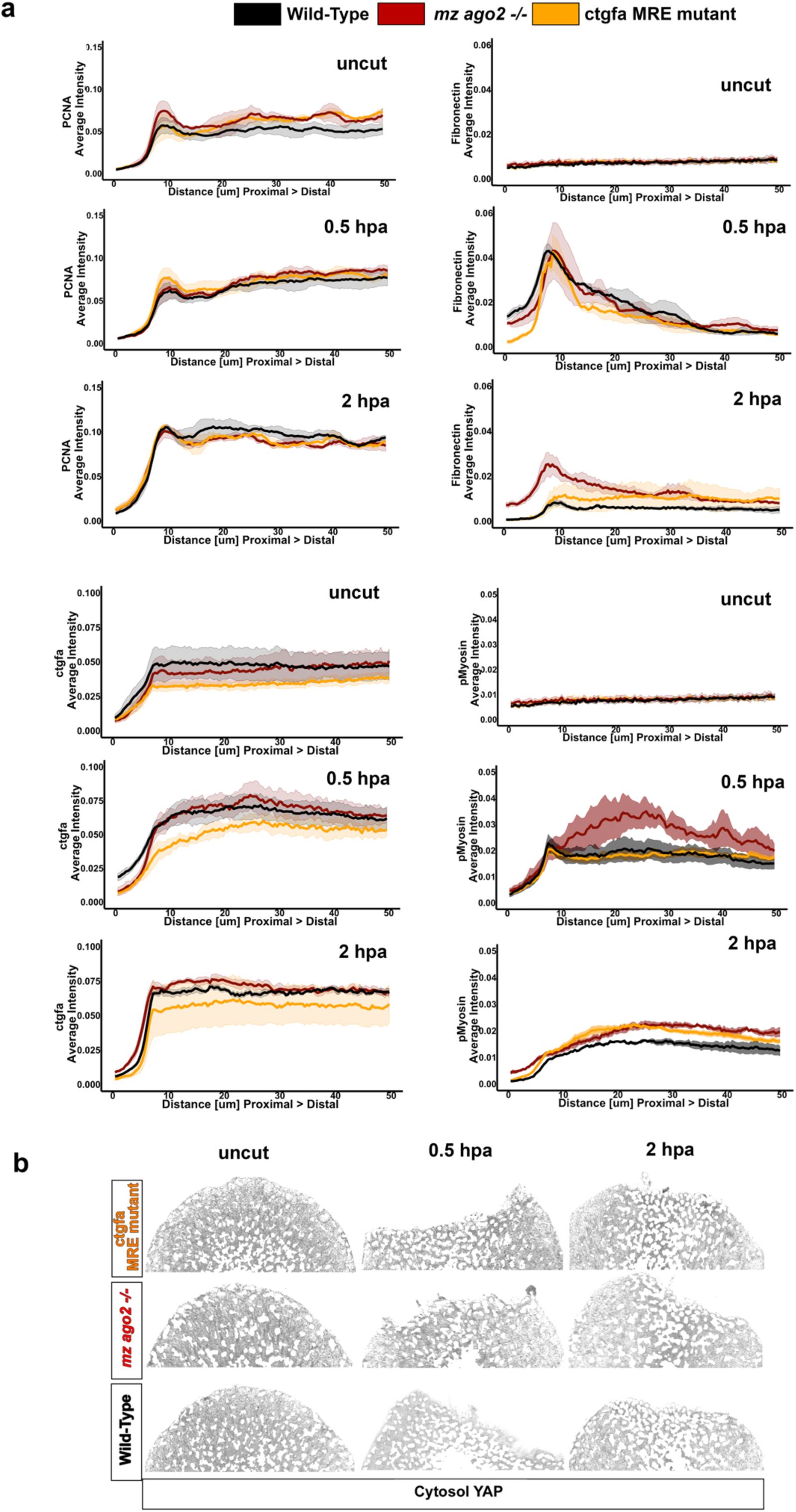
Spatial profiles of CAM proteins and YAP Nuclear/Cytosol quantification. (a) Fluorescent intensity profiles for the indicated proteins and treatments depicting the protein quantification per zebrafish fin fold represented in Fig.5 a-c and Supplementary Fig. 7 a-c). Pixel matrix of intensity was reconstructed for 50 μm within the wound edge to the fish body. Pixel mean was calculated and normalized for the maximum value to obtain pixel intensity profile. The intensity profile of 4-6 fish was combined and the smooth distribution of intensity was calculated. Solid lines show the mean for each genotype, while the grey ribbons show standard error. **(b)** Representative confocal lateral images of YAP immunostaining in the uncut fin fold. A DAPI binary mask was used to generate the Nuclear YAP (Fig. 5 c) and Cytosol YAP images.

**Supplementary Table 1. AGO2-peaks localization.** The table shows the chromosome location and strand, gene name and Ensembl Gene ID and the peak p-value calculated with the Piranha (Methods). The last two columns show the number of genes associated with AGO2 and the number of peaks identified from the AGO2-HITS-CLIP.

**Supplementary Table 2. AGO2-miRNAs are clustered in 155 families.** The table lists the miRNAs associate with AGO2 complex in endothelial cells (HUVEC and HUAEC). For each miRNA, the mature sequence and the miRbase ID are shown. MiRNAs are grouped in families based on the 7-mer SEED region (column Family and SEED).

**Supplementary Table 3. miRNA-CAM-MRE interaction network.** The table represents the interaction network between the selected CAM-MRE and the endothelial miRNA within the AGO2 complex. The network was generated using TargetScan v. 7.0 (method) using CAM peaks listed in Table 1 and miRNA family listed in Table 2. Site_type, UTR_start and UTR_end columns represent the seed interaction type base (6mer < 7mer-1a < 7mer-m8 < 8mer-1a) and the localization of the interaction site within the Sensor_MRE_sequence (Table 4).

**Supplementary Table 4. CAM-MRE sequences.** The table lists the selected CAM sequence used for the Sensor-seq assay (method) and reported in Fig. 2. Canonical Gene name, MRE identity, extended MRE sequence, and length of the sequences cloned into the Sensor-Seq vector (method) are reported.

**Supplementary Table 5. Proteomics results.** The table lists the proteomics analysis performed on HUVEC cells seed on soft (3 kPa) or stiff (30 kPa) PDMS gel, or infected with AGO2 gRNA or not-target gRNA. The fold change values were used to generate the scatter plot in Figure 3b.

**Supplementary Table 6. CAM-MRE gRNA.** The table lists the selected CAM-MRE sequence, and the gRNA used for destabilizing the interaction with the miRNA (method)

## Acknowledgements

We thank Meredith Cavanaugh for help with zebrafish husbandry. We thank the Wellcome Trust Centre for Cell-Matrix Research for technical support with tissue culture assays. We thank Melanie Trombly and Angela Anderson for proofreading the manuscript, Jay Humphrey and Valentina Greco for critical reading. We thank Brian Coon for assistance during the preparation of the CRISR/Cas9 experiment. We thank Dr. Jun Lun’s laboratory for providing miRNA reporter lentiviral plasmid. JS was funded by a Biotechnology and Biological Sciences Research Council (BBSRC) David Phillips Fellowship (BB/L024551/1). VM was partially supported by a studentship from the Sir Richard Stapley Educational Trust. Mass spectrometry was carried out at the Wellcome Trust Centre for Cell-Matrix Research (WTCCMR; 203128/Z/16/Z) Proteomics Core Facility. This work was supported by USPHS grant RO1 GM47214 to MAS and RO1 HL130246 to SN.

## Author contributions

SN and MS conceived the project. AM and TD performed experiments, analyzed data and AM developed the computational data analysis. WA performed and analyzed zebrafish fin fold regeneration experiments. LB performed and analyzed the experiments in Supplementary Fig 2b and c, Supplementary figure 3b-d and 6. SJA performed cell culture experiments in Supplementary Fig.3a. NB and CJ performed the 3D cell-culture experiments and analyzed the data. DL, MG, JZ, and MG develop the sequence data processing, analysis (mapping, peak calling) of the AGO2-HITS-CLIP experiment. VM and JS performed mass spectrometry proteomic analysis. DMK made the miRNA and RNA-sequencing libraries. AM, TD, MS and SN designed the experiments. MS and SN wrote the manuscript. All the authors edited the manuscript.

## References

1. Cyron, C.J. & Humphrey, J.D. Growth and Remodeling of Load-Bearing Biological Soft Tissues. Meccanica 52, 645–664 (2017).

2. Gilbert, P.M.& Weaver, V.M. Cellular adaptation to biomechanical stress across length scales in tissue homeostasis and disease. Semin Cell Dev Biol 67, 141–152 (2017).

3. Humphrey, J.D., Dufresne, E.R.& Schwartz, M.A. Mechanotransduction and extracellular matrix homeostasis. Nat Rev Mol Cell Biol 15, 802–12 (2014).

4. Humphrey, J.D. Vascular adaptation and mechanical homeostasis at tissue, cellular, and sub-cellular levels. Cell Biochem Biophys 50, 53–78 (2008).

5. Seki, E.& Brenner, D.A. Recent advancement of molecular mechanisms of liver fibrosis. J Hepatobiliary Pancreat Sci 22, 512–8 (2015).

6. Huang, S.& Ingber, D.E. Cell tension, matrix mechanics, and cancer development. Cancer Cell 8, 175–6 (2005).

7. Sun, Z., Guo, S.S.& Fassler, R. Integrin-mediated mechanotransduction. J Cell Biol 215, 445–456 (2016).

8. Bartel, D.P. MicroRNAs: target recognition and regulatory functions. Cell 136, 215–33 (2009).

9. Ebert, M.S.& Sharp, P.A. Roles for microRNAs in conferring robustness to biological processes. Cell 149, 515–24 (2012).

10. Guo, H., Ingolia, N.T., Weissman, J.S.& Bartel, D.P. Mammalian microRNAs predominantly act to decrease target mRNA levels. Nature 466, 835–40 (2010).

11. Herranz, H.& Cohen, S.M. MicroRNAs and gene regulatory networks: managing the impact of noise in biological systems. Genes Dev 24, 1339–44 (2010).

12. Tsang, J., Zhu, J.& van Oudenaarden, A. MicroRNA-mediated feedback and feedforward loops are recurrent network motifs in mammals. Mol Cell 26, 753–67 (2007).

13. Kasper, D.M.et al. MicroRNAs Establish Uniform Traits during the Architecture of Vertebrate Embryos. Dev Cell 40, 552–565 e5 (2017).

14. Olson, E.N. MicroRNAs as therapeutic targets and biomarkers of cardiovascular disease. Sci Transl Med 6, 239ps3 (2014).

15. Nicoli, S., Knyphausen, C.P., Zhu, L.J., Lakshmanan, A.& Lawson, N.D. miR-221 is required for endothelial tip cell behaviors during vascular development. Dev Cell 22, 418–29 (2012).

16. Nicoli, S.et al. MicroRNA-mediated integration of haemodynamics and Vegf signalling during angiogenesis. Nature 464, 1196–200 (2010).

17. Ristori, E.et al. A Dicer-miR-107 Interaction Regulates Biogenesis of Specific miRNAs Crucial for Neurogenesis. Dev Cell 32, 546–60 (2015).

18. Bracken, C.P., Scott, H.S.& Goodall, G.J. Anetwork-biology perspective of microRNA function and dysfunction in cancer. Nat Rev Genet 17, 719–732 (2016).

19. Pelaez, N.& Carthew, R.W. Biological robustness and the role of microRNAs: a network perspective. Curr Top Dev Biol 99, 237–55 (2012).

20. Pasquinelli, A.E. MicroRNAs and their targets: recognition, regulation and an emerging reciprocal relationship. NatRev Genet 13, 271–82 (2012).

21. Hafner, M.et al. Transcriptome-wide identification of RNA-binding protein and microRNA target sites by PAR-CLIP. Cell 141, 129–41 (2010).

22. Chi, S.W., Zang, J.B., Mele, A.& Darnell, R.B. ArgonauteHITS-CLIP decodes microRNA-mRNA interaction maps. Nature 460, 479–86 (2009).

23. Byfield, F.J., Reen, R.K., Shentu, T.P., Levitan, I.& Gooch, K.J. Endothelialactin and cell stiffness is modulated by substrate stiffness in 2D and 3D. J Biomech 42, 1114–9 (2009).

24. Grimson, A.et al.MicroRNA targeting specificity in mammals: determinants beyond seed pairing. Mol Cell 27, 91–105 (2007).

25. Saphirstein, R.J.& Morgan, K.G. The contribution of vascular smooth muscle to aortic stiffness across length scales. Microcirculation 21, 201–7 (2014).

26. Mullokandov, G.et al. High-throughput assessment of microRNA activity and function using microRNA sensor and decoy libraries. Nat Methods 9, 840–6 (2012).

27. Kamata, M., Liang, M., Liu, S., Nagaoka, Y.& Chen, I.S. Livecell monitoring of hiPSC generation and differentiation using differential expression of endogenous microRNAs. PLoS One 5, e11834 (2010).

28. Discher, D.E., Janmey, P.& Wang, Y.L. Tissue cells feel and respond to the stiffness of their substrate. Science 310, 1139–43 (2005).

29. Kim, Y.K., Kim, B.& Kim, V.N. Re-evaluation of the roles of DROSHA, Export in 5, and DICER in microRNA biogenesis. Proc Natl Acad Sci U S A 113, E1881–9 (2016).

30. Dupont, S.et al. Role of YAP/TAZ in mechanotransduction. Nature 474, 179–83 (2011).

31. Kumar, A.et al. Talin tension sensor reveals novel features of focal adhesion force transmission and mechanosensitivity. J Cell Biol 213, 371–83 (2016).

32. Bassett, A.R.et al. Understanding functional miRNA-target interactions in vivo by site-specific genome engineering. Nat Commun 5, 4640 (2014).

33. hi-Wen, X., Leask, A.& Abraham, D. Regulationand function of connective tissue growth factor/CCN2 in tissue repair, scarring and fibrosis. Cytokine Growth Factor Rev 19, 133–44 (2008).

34. Mateus, R.et al. Control of tissue growth by Yap relies on cell density and F-actin in zebrafish fin regeneration. Development 142, 2752–63 (2015).

35. Kawakami, A., Fukazawa, T.& Takeda, H. Earlyfin primordia of zebrafish larvae regenerate by a similar growth control mechanism with adult regeneration. DevDyn 231, 693–9 (2004).

36. Mathew, L.K.et al. Comparative expression profiling reveals an essential role for raldh2 in epimorphic regeneration. J Biol Chem 284, 33642–53 (2009).

37. Mateus, R.et al. In vivo cell and tissue dynamics underlying zebrafish fin fold regeneration. PLoS One 7, e51766 (2012).

38. Cifuentes, D.et al. A novel miRNA processing pathway independent of Dicer requires Argonaute2 catalytic activity. Science 328, 1694–8 (2010).

39. Amelio, I.et al. miR-24 triggers epidermal differentiation by controlling actin adhesion and cell migration. J Cell Biol 199, 347–63 (2012).

40. Nechiporuk, A.& Keating, M.T. Aproliferation gradient between proximal and msxb-expressing distal blastema directs zebrafish fin regeneration. Development 129, 2607–17 (2002).

41. Mori, M.et al. Hippo signaling regulates microprocessor and links cell-density-dependent miRNA biogenesis to cancer. Cell 156, 893–906 (2014).

42. Chaulk, S.G., Lattanzi, V.J., Hiemer, S.E., Fahlman, R.P.& Varelas, X. TheHippo pathway effectors TAZ/YAP regulate dicer expression and microRNA biogenesis through Let-7. J Biol Chem 289, 1886–91 (2014).

43. Davis, B.N., Hilyard, A.C., Lagna, G.& Hata, A. SMADproteins control DROSHA-mediated microRNA maturation. Nature 454, 56–61 (2008).

44. Felix, M.A.& Wagner, A. Robustness and evolution: concepts, insights and challenges from a developmental model system. Heredity (Edinb) 100, 132–40 (2008).

45. Mouw, J.K.et al. Tissue mechanics modulate microRNA-dependent PTEN expression to regulate malignant progression. Nat Med 20, 360–7 (2014).

46. Liu, G.et al.miR-21 mediates fibrogenic activation of pulmonary fibroblasts and lung fibrosis. J Exp Med 207, 1589–97 (2010).

47. Cushing, L.et al. miR-29 is a major regulator of genes associated with pulmonary fibrosis. Am J Respir Cell Mol Biol 45, 287–94 (2011).

48. Herrera, J.et al.Dicer1 Deficiency in the IPF Fibroblastic Focus Promotes Fibrosis by Suppressing MicroRNA Biogenesis. Am J Respir Crit Care Med (2018).

49. Parker, M.W.et al.Fibrotic extracellular matrix activates a profibrotic positive feedback loop. J Clin Invest 124, 1622–35 (2014).

50. Pandit, K.V.& Milosevic, J. MicroRNAregulatory networks in idiopathic pulmonary fibrosis. Biochem Cell Biol 93, 129–37 (2015).

51. Wynn, T.A.& Ramalingam, T.R. Mechanismsof fibrosis: therapeutic translation for fibrotic disease. Nat Med 18, 1028–1040 (2012).

52. Uren, P.J.et al.Site identification in high-throughput RNA-protein interaction data. Bioinformatics 28, 3013–20 (2012).

53. Dobin, A.et al. STAR: ultrafast universal RNA-seq aligner. Bioinformatics 29, 15–21 (2013).

54. Li, H.et al. The Sequence Alignment/Map format and SAMtools. Bioinformatics 25, 2078–9 (2009).

55. Kozomara, A.& Griffiths-Jones, S. miRBase: annotating high confidence microRNAs using deep sequencing data. Nucleic Acids Res 42, D68–73 (2014).

56. Agarwal, V., Bell, G.W., Nam, J.W.& Bartel, D.P. Predictingeffective microRNA target sites in mammalian mRNAs. Elife 4 (2015).

57. Aranguren, X.L.et al.Unraveling a novel transcription factor code determining the human arterial-specific endothelial cell signature. Blood 122, 3982–92 (2013).

58. Ritchie, M.E.et al.limma powers differential expression analyses for RNA-sequencing and microarray studies. Nucleic Acids Res 43, e47 (2015).

59. Narayanan, A.et al.In vivo mutagenesis of miRNA gene families using a scalable multiplexed CRISPR/Cas9 nuclease system. Sci Rep 6, 32386 (2016).

60. Bodenhofer, U., Bonatesta, E., Horejs-Kainrath, C. & Hochreiter, S. msa: an R package for multiple sequence alignment. Bioinformatics 31, 3997–9 (2015).

61. Elosegui-Artola, A.et al. Mechanical regulation of a molecular clutch defines force transmission and transduction in response to matrix rigidity. Nat Cell Biol 18, 540–8 (2016).

62. Robinson, M.D., McCarthy, D.J.& Smyth, G.K. edgeR: a Bioconductor package for differential expression analysis of digital gene expression data. Bioinformatics 26, 139–40 (2010).

63. McCarthy, D.J., Chen, Y.& Smyth, G.K. Differentialexpression analysis of multifactor RNA-Seq experiments with respect to biological variation. Nucleic Acids Res 40, 4288–97 (2012).

64. Goeminne, L.J., Gevaert, K.& Clement, L. Peptide-level Robust Ridge Regression Improves Estimation, Sensitivity, and Specificity in Data-dependent Quantitative Label-free Shotgun Proteomics. Mol Cell Proteomics 15, 657–68 (2016).

65. Tyanova, S.et al. The Perseus computational platform for comprehensive analysis of (prote)omics data. Nat Methods 13, 731–40 (2016).

66. Smyth, G.K. Linearmodels and empirical bayes methods for assessing differential expression in microarray experiments. Stat Appl Genet Mol Biol 3, Article3 (2004).

67. Berginski, M.E.& Gomez, S.M. The Focal Adhesion Analysis Server: a web tool for analyzing focal adhesion dynamics. F1000Res 2, 68 (2013).

68. Gutierrez, E.& Groisman, A. Measurements of elastic moduli of silicone gel substrates with a microfluidic device. PLoS One 6, e25534 (2011).

69. Han, S.J., Oak, Y., Groisman, A.& Danuser, G. Tractionmicroscopy to identify force modulation in subresolution adhesions. Nat Methods 12, 653–6 (2015).

70. Kapacee, Z.et al. Tension is required for fibripositor formation. Matrix Biol 27, 371–5 (2008).

71. Schindelin, J.et al. Fiji: an open-source platform for biological-image analysis. Nat Methods 9, 676–82 (2012).

72. Le Guyader, D.et al. Origins and unconventional behavior of neutrophils in developing zebrafish. Blood 111, 132–41 (2008).

73. Sneddon, I.N. The relation between load and penetration in the axisymmetric boussinesq problem for a punch of arbitrary profile. Int. J. Engng Sci. 3, 47–57 (1965).

74. McCall, M.N.et al. MicroRNA profiling of diverse endothelial cell types. BMC Med Genomics 4, 78 (2011).

75. Krzywinski, M.et al. Circos: an information aesthetic for comparative genomics. Genome Res 19, 1639–45 (2009).

